# Biomechanical interactions of *Schistosoma mansoni* eggs with vascular endothelial cells facilitate egg extravasation

**DOI:** 10.1101/2021.09.16.459846

**Authors:** Yi-Ting Yeh, Danielle E. Skinner, Ernesto Criado-Hidalgo, Natalie Shee Chen, Antoni Garcia-De Herreros, Nelly El-Sakary, Lawrence Liu, Shun Zhang, Shu Chien, Juan C. Lasheras, Juan C. del Álamo, Conor R. Caffrey

**Affiliations:** Department of Mechanical and Aerospace Engineering, University of California, San Diego, 9500 Gilman Drive, La Jolla, California, United States; Department of Bioengineering, University of California, San Diego, 9500 Gilman Drive, La Jolla, California, United States; Institute of Engineering in Medicine, University of California, San Diego, 9500 Gilman Drive, La Jolla, California, United States; Center for Discovery and Innovation in Parasitic Diseases (CDIPD), Skaggs School of Pharmacy and Pharmaceutical Sciences, University of California, San Diego, 9500 Gilman Drive, La Jolla, California, United States; Department of Mechanical Engineering, University of Washington, 3900 E Stevens Way NE, Seattle, Washington, United States; Center for Cardiovascular Biology, University of Washington, 850 Republican St, Seattle Washington, United States; Institute for Stem Cell and Regenerative Medicine, University of Washington, 850 Republican St, Seattle Washington, United States

## Abstract

The eggs of the parasitic blood fluke, *Schistosoma*, are the main drivers of the chronic pathologies associated with schistosomiasis, a disease of poverty afflicting approximately 220 million people worldwide. Eggs laid by *Schistosoma mansoni* in the bloodstream of the host are encapsulated by vascular endothelial cells (VECs), the first step in the migration of the egg from the blood stream into the lumen of the gut and eventual exit from the body. The biomechanics associated with encapsulation and extravasation of the egg are poorly understood. We demonstrate that *S. mansoni* eggs induce VECs to form two types of membrane extensions during encapsulation; filopodia that probe eggshell surfaces and intercellular nanotubes that presumably facilitate VEC communication. Encapsulation efficiency, the number of filopodia and intercellular nanotubes, and the length of these structures depend on the egg’s vitality and, to a lesser degree, its maturation state. During encapsulation, live eggs induce VEC contractility and membranous structures formation, in a Rho/ROCK pathway-dependent manner. Using elastic hydrogels embedded with fluorescent microbeads as substrates to culture VECs, live eggs induce VECs to exert significantly greater contractile forces during encapsulation than dead eggs, which leads to 3D deformations on both the VEC monolayer and the flexible substrate underneath. These significant mechanical deformations cause the VEC monolayer tension to fluctuate with eventual rupture of VEC junctions, thus facilitating egg transit out of the blood vessel. Overall, our data on the mechanical interplay between host VECs and the schistosome egg improve our understanding of how this parasite manipulates its immediate environment to maintain disease transmission.

## Introduction

Schistosomiasis is caused by several species of the *Schistosoma* blood fluke. Transmitted by freshwater snails, the disease is prevalent in sub-Saharan Africa and parts of Southeast Asia and South America, with approximately 220 million people infected (1). During infection with *Schistosoma mansoni*, paired male and female worms wander through the mesenteric and hepatic portal venous systems that drain the alimentary canal. Females lay hundreds of eggs per day, and these come into direct contact with vascular endothelial cells (VECs) (2–4). VECs first migrate over and encapsulate the eggs, initiating an inflammatory granulomatous response that facilitates egg extravasation. Egg extravasation is critical for subsequent granuloma-mediated egg transport through the gut wall into the lumen, and eventually, the release of eggs into the external environment with the feces (3, 5–8). Eggs that fail to extravasate are swept away by the blood flow and become trapped in the liver where they induce granulomata and eventually fibrosis which results in pain and malaise with often life-threatening sequela over the course of years or even decades. (9–11).

During and after their interaction with VECs, eggs grow and mature (12), and maturation is distinguished by the differentiation of various organs and tissues in the developing miracidium that is contained within the eggshell (13–15). The eggshell is composed primarily of chitin and has a complex surface topography with several structures that could interact with VECs (16, 17). One of the eggshell’s most prominent features is an evenly spaced pattern of microspine structures with an average length of 200-300 nm and a diameter of 60 nm (18–21). The eggshell also exhibits nanometer-size pores through which the developing embryo releases egg secretory proteins (ESPs) (12, 22–24). These secreted molecules induce (i) the adhesion of VECs by increasing VEC expression of the adhesion molecules, ICAM-1 and VCAM-1, and (ii) VEC proliferation that facilitates egg-encapsulation and the subsequent recruitment of host leukocytes to initiate granuloma formation (7, 25).

Extravasation of the schistosome egg requires mechanical forces to push the egg in the direction of the extravascular space and bring it into direct contact with the blood vessel’s basement membrane. However, the rigidity of the shell limits the transmission of appreciable forces to its surroundings (3, 4, 26). Muscular contractions of the female worm and/or migration of the host’s VECs over the egg could generate the forces necessary for egg transmigration (27). Although the movement forces of the male worm have been quantified (28), there are no measurements of female-generated worm forces, including those during egg-deposition. VEC actomyosin contractility, coupled with substrate adhesion, create tensile forces that are transmitted between neighboring cells via adherens junctions, and, together, these contribute to the generation of mechanical tension at the tissue level (29). Tensional homeostasis within VEC monolayers plays a role in a wide range of processes such as shear stress mechanosensing (30), leukocyte trafficking (31) and VEC-pathogen interactions (32, 33). However, the VEC response to schistosome egg contact, and specifically whether and how VECs remodel their actomyosin cytoskeleton and actively generate mechanical forces to drive the encapsulation of eggs, are poorly understood. Successful migration of the schistosome egg into the lumen of the bowel is crucial to minimizing the accumulative and, often, life-threatening pathologies that are associated with those eggs that become trapped (3). Thus, investigating the biomechanics of egg extravasation could help identify the processes and molecules that facilitate egg excretion, thereby offering a better understanding of disease pathogenesis.

Here, we first analyzed the ultrastructure of *S. mansoni* eggs and VECs during the encapsulation process and identified two types of VEC membrane extensions: filopodia and intercellular nanotubes (NTs). We showed that the number and length of filopodia and NTs during encapsulation of eggs are influenced by the age and vitality of the parasite embryo. We also performed three-dimensional traction force microscopy (3D-TFM) measurements on cultured VEC monolayers to show that live, but not dead, eggs increased VEC contractility, which resulted in eggs being pushed against the substrate to eventually rupture the VEC intercellular junctions. Together, our data demonstrate a series of intricate and coordinated biomechanical events that facilitate egg extravasation into the underlying host tissues.

## Results

### VECs form actin-rich filopodia structures that physically probe eggshell microspines during encapsulation of *S. mansoni* eggs

To investigate how VECs encapsulate *S. mansoni* eggs, we placed mature eggs onto VEC monolayers cultured *in vitro*. After 24 h, the VECs had fully encapsulated the eggs as evidenced by immunostaining of the endothelial junction protein, VE-cadherin, which was detected in all z-planes of view (basal, middle and apical; Fig. 1A). We also imaged the interactions between mature eggs and VECs using scanning electron microscopy (SEM) at 4 and 24 h (Fig. 1B). At 4 h, VECs protruded membrane extensions called filopodia, to physically interact with the eggs; by 24 h, the VECs had completely covered the eggs. Following the release of eggs by female worms, immature eggs interact with VECs and they become mature over the course of several days (13–15). Immature eggs are about two-thirds the length and width of mature eggs (SI table 1). DAPI staining showed that most of the nuclei in immature eggs were evenly dispersed throughout the embryo, whereas those in more mature eggs have generated a well-defined circular neural mass in the center of the developing miracidium (SI Fig. 1B and SI Table 1). Furthermore, SEM showed that the surface of the eggshell was covered by microspine structures (Fig. 1C), as previously reported (21). By comparing mature and immature eggs, we found that similar microspines were present on both with an average length and width of ~200 nm (Figs. 1C and D), although the microspine shapes on immature eggs were more irregular (Fig. 1C). VECs developed long filopodia that intimately contacted the egg microspines (Fig. 1E). Using confocal microscopy and phalloidin immunostaining, we confirmed the presence of F-actin in these VEC filopodia (Fig. 1F). In particular, F-actin-rich filopodia were observed on the eggshell’s apical end (Fig. 1F). The filopodia were integrated into the F-actin cytoskeleton of the VECs, as evidenced by tracking these structures in subsequent z-planes (SI Fig. 2 and SI Movie 1).

**Figure 1.**
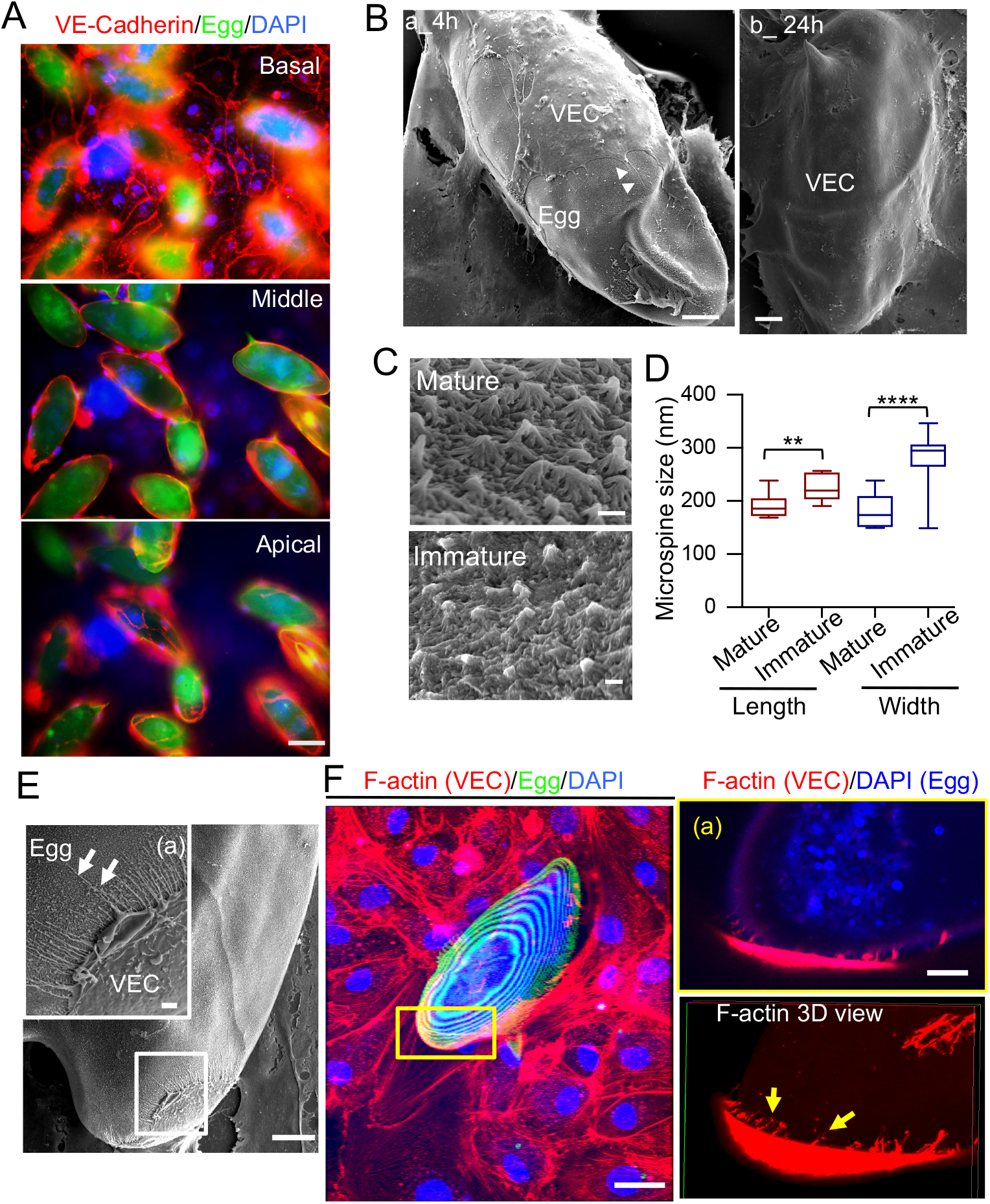
VECs generate filopodia while encapsulating *S. mansoni* eggs. **(A)** Immunostaining shows VECs (VE-Cadherin, red) on the basal, middle and apical areas of live *S. mansoni* mature eggs, which are auto-fluorescently green. DAPI was used to stain for cell nuclei. Scale bar: 50 μm. **(B)** SEM images demonstrating the interaction between VECs and the eggshell: (a) At 4 h, part of an eggshell was covered by VECs (scale bar: 10 μm), Arrowheads point VEC membrane protrusions on the eggshell; (b) at 24 h, the eggshell was fully covered by VECs (scale bar: 10 μm). **(C)** Image of microspines in mature and immature eggshells (scale bar: 0.2 μm). **(D)** Quantification of the length and width of microspines in mature and immature eggshells. ** and **** indicate respectively p < 0.01 and p < 0.0001 by Student’s *t-*test; 12-15 microspines were quantified in each condition. **(E)** SEM image of the interaction between the *S. mansoni* egg and VEC filopodia after 4 h (scale bar = 10 μm); (a) enlarged image of the boxed region of interest showing VEC filopodia (arrows) on the eggshell surface. Scale bar: 1 μm. **(F)** Confocal microscopy images of actin filaments. VECs and eggs were fixed, and immunostained with rhodamine phalloidin (red). Scale bar: 20 μm. (a) Enlarged image of the boxed region of interest (showing the presence of VEC actin-rich filopodia during VEC encapsulation of the egg. Scale bar: 5 μm. Lower panel is the 3D view and arrows indicate actin filaments on the eggshell surface.

### In addition to filopodia, VECs generate intercellular nanotubes during the encapsulation of *S. mansoni* eggs

After 4 h of co-culture of mature eggs with VECs, SEM revealed the presence of VEC intercellular nanotubes (NTs) in addition to filopodia on *S. mansoni* eggs. NTs are specialized membrane extensions that connect neighboring cells (34, 35) and mediate cell-cell communications by internal transport of signaling molecules such as proteins, lipids and nucleic acids (36). During encapsulation of the egg, VECs were observed to elaborate NTs from their training edges which maintained connections with the basal endothelium (Fig. 2A). At the same time, filopodia were observed at the advancing front of the VECs exploring the egg surface (Figs. 2A–B).

**Figure 2.**
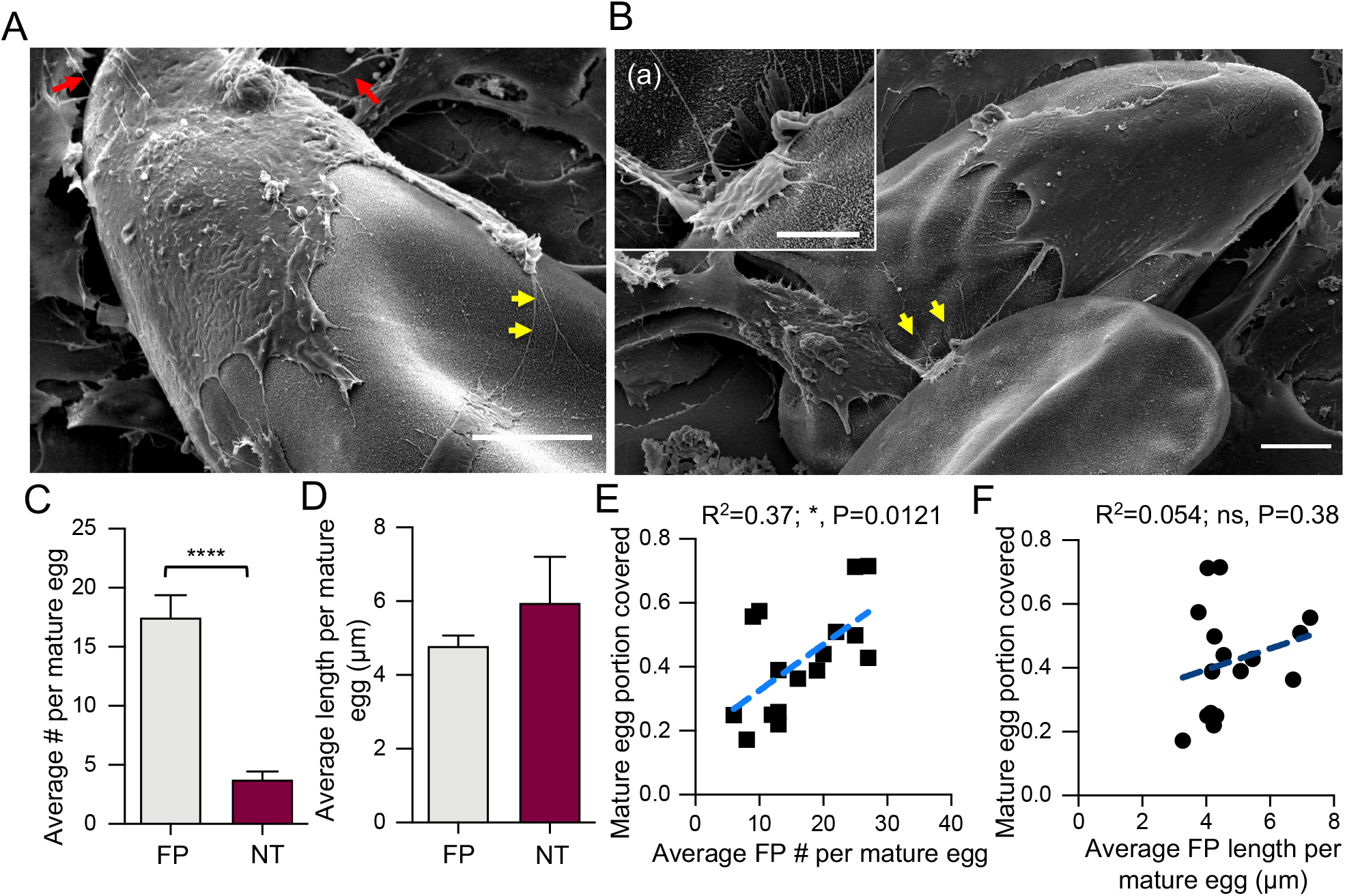
VECs interacting with *S. mansoni* eggs display two types of membrane protrusions. **(A)** During encapsulation, VECs extend filopodia (FP; yellow arrows), to probe the egg surface, and intercellular nanotubes (NTs; red arrows) to connect with neighboring cells (scale bar: 10 μm). **(B)** Representative SEM image to illustrate how filopodial probing facilitates VEC migration over eggs: (a) enlarged image of the probing FP corresponding to the region indicated by the yellow arrowheads (scale bar: 4 μm). **(C)** Average number of FPs and NTs formed per mature egg after 4 h. **(D)** Average length of FPs and NTs formed per mature egg after 4 h. Bar plots in **C** and **D** represent mean ± s.e.m. **** indicate p < 0.0001 by Student’s *t*-test. **(E)** Egg portion covered by VECs as a function of average FP number. **(F)** Egg portion covered by VECs as a function of average FP length. For both **E** and **F**, * indicates statistically significant Pearson correlation (p = 0.0121). For **C** - **F**, a total of 16 mature eggs from three independent experiments were analyzed.

The average numbers of VEC filopodia and intercellular NTs per egg after a 4 h interaction were approximately 17 and 4, respectively (being significantly different from each other at p < 0.0001), although the average length of filopodia and NTs per egg was not significantly different (p > 0.05), being 4.78 ± 0.28 μm and 5.96 ± 1.24 μm, respectively (Figs. 2C–D). Next, we analyzed whether VEC filopodia number and length correlated with the portion of the egg that was covered by VECs. There was a modest correlation for filopodia number (R^2^ = 0.37, p = 0.01, Fig. 2E), but there was no correlation for filopodia length (R^2^ = 0.054, p = 0.38; Fig. 2F). Together, these data demonstrate that there are two distinct types of VEC membrane protrusions, *i.e*., filopodia and NTs, during the egg encapsulation process, and that there is a positive correlation between VEC filopodia number and the cells’ ability to encapsulate eggs. The data may suggest that during the egg-VEC interaction, VECs favor exploring their environment over communicating with neighboring cells by generating more forward-projecting filopodia than rear-facing intercellular NTs.

### Live eggs constitute a greater stimulus than dead eggs to the encapsulation process by VECs

To understand whether the induction of VEC filopodia was influenced by the embryo’s developmental state and vitality, we tested the response of VECs to live immature and mature eggs, as well as mature eggs that had been killed with sodium azide (hereafter referred to as dead eggs). The vitality of the live mature eggs was confirmed by their ability to hatch in water under a bright light for 40 min. Hatching efficiency was ~80% (SI Fig. 1A); also, we observed that those eggs capable of hatching always contained a moving miracidium (SI Movie 2). SEM imaging showed that the average number of filopodia and NTs per egg depended on egg vitality (~2.5-fold more in live eggs, p = 0.0002, mature *vs.* dead; p < 0.0001, immature *vs.* dead) but not developmental state after a 4 h incubation with VECs (Figs. 3A–B). Immature eggs induced VECs to form significantly (~2-fold) longer filopodia than either mature (p = 0.0002) or dead eggs (p < 0.0001). For NTs, length was a function of egg vitality (~5-fold longer in both mature (p = 0.041) and immature eggs (p = 0.01), *vs.* dead eggs; Fig. 3C) but not development.

**Figure 3.**
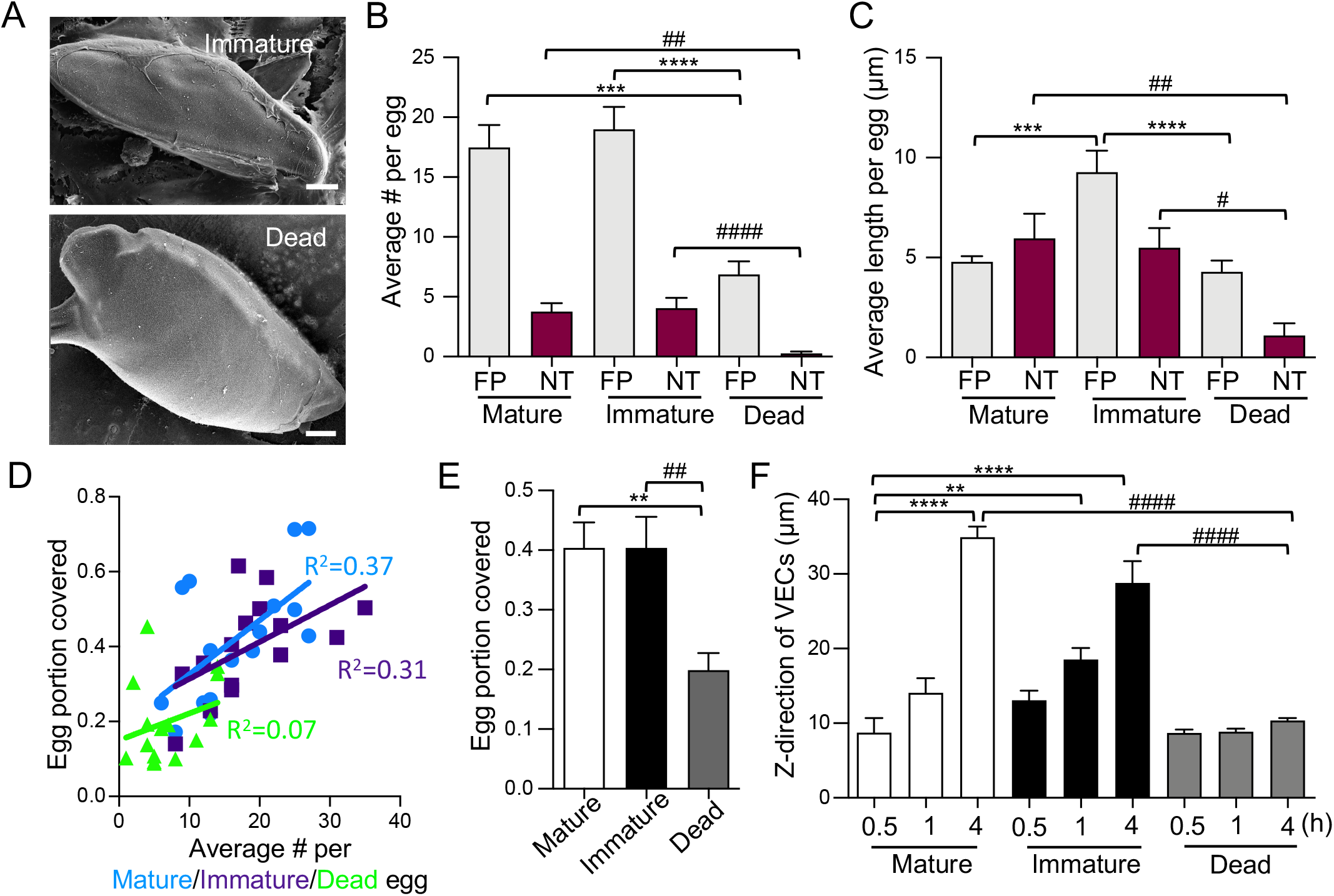
Encapsulation of *S. mansoni* eggs by VEC filopodia depends on egg vitality. **(A)** SEM images of VECs encapsulating a live immature egg and a dead mature egg after 4 h (scale bar: 10 μm). **(B)** Number of FPs and NTs per egg (mean ± s.e.m.). *** and **** indicate p < 0.001 and p < 0.0001 when comparing FPs. ## and #### indicate p < 0.01 and p < 0.0001 when comparing NTs. **(C)** Length of FP and NTs per egg (mean ± s.e.m.). *** and **** indicate p < 0.001 and p < 0.0001 when comparing FPs. # and ## indicate p < 0.05 and p<0.01 when comparing NTs. For **B** and **C**, a one-way ANOVA with Tukey’s post-test was used. **(D)** The portion of eggs covered by VECs as a function of the average number of FPs per egg at 4 h. Mature (light blue circles) and immature live eggs (purple squares) show significant Pearson correlations which p value = 0.012 and 0.029, respectively), whereas dead eggs do not (p = 0.075). **(E)** The portion of live mature (white), immature (black) and dead (gray) eggs covered by VECs after 4 h. ** indicates p < 0.01 and ## indicates p < 0.01 using a one-way ANOVA with Tukey’s post-test. For **A** – **E**, a total of 16, 15 and 14 live mature, live immature and dead eggs from three independent experiments were analyzed. **(F)** Quantification of the encapsulation dynamics as a function of the maximum height (z-direction; mean ± s.e.m.) reached by VECs at different time points. ** and **** indicate p < 0.01and p < 0.0001, respectively, using a one-way ANOVA with Dunnett’s post-test. #### indicates p < 0.0001 when comparing 4 h mature, immature and dead eggs, using a one-way ANOVA with Tukey’s post-test. Across three independent experiments, 7-12 eggs were analyzed for each condition.

Next, we investigated whether, after 4 h, there was a correlation between the number of VEC filopodia per egg and the portion of the egg covered by VECs, for each of the three egg preparations (Fig. 3D). There was a modest correlation for live mature (R^2^ = 0.37, p = 0.012) and immature eggs (R^2^ = 0.31, p = 0.039), but there was no significant correlation for dead eggs (R^2^ = 0.07, p = 0.075). Also, the average egg portion covered by VECs was greater for both mature (0.4; p = 0.0025) and immature eggs (0.2; p = 0.0048) *vs*. dead eggs (Fig. 3E). To understand the dynamics of encapsulation, co-cultures of mature and immature eggs, and VECs, were fixed at 0.5, 1 and 4 h, and imaged by confocal microscopy (SI Fig. 3). VECs migrated significantly faster (~3-fold) over both live immature and mature eggs compared to dead eggs (p < 0.0001 for both mature and immature eggs *vs.* dead eggs at 4 h; Fig. 3F) and due, perhaps, to immature eggs being smaller, full encapsulation had already occurred by 4 h (Fig. 3F and SI Fig. 3). These data indicate that live eggs induce filopodia and NTs during encapsulation, whereas dead eggs provide less of a stimulus to the encapsulation process.

### Rho/ROCK-mediated VEC contractility is involved in the formation of filopodia and NTs, and the encapsulation of eggs

Having established that live eggs induce VEC filopodia and NTs during encapsulation, we investigated the mechanisms that may regulate VEC motility. Phosphorylation of the myosin regulatory light chain (MLC) is known to elicit the contraction of the actin cytoskeleton that facilitates cell motility (37). Accordingly, we immunostained VECs for phosphorylation of MLC after a 4 h incubation with live mature and immature *S. mansoni* eggs. Confocal microscopy showed that active phosphor-MLC was more densely located in those VECs that were in direct contact or closely associated with eggs in the basolateral focal plane of the VEC monolayers (Figs 4A, B). This pattern of increased phosphor-MLC in VECs was similar for both live mature and immature eggs. Pre-treating VECs with 20 μM Y-27632 or ML-7, small molecule inhibitors of Rho/ROCK and MLC kinase, respectively, significantly decreased the phosphor-MLC signal upon incubation with the live mature eggs (Fig. 4C). The data demonstrate that live eggs induce VEC myosin activity and suggest that local myosin activity enables VEC contractility at egg contact sites to facilitate encapsulation.

**Figure 4.**
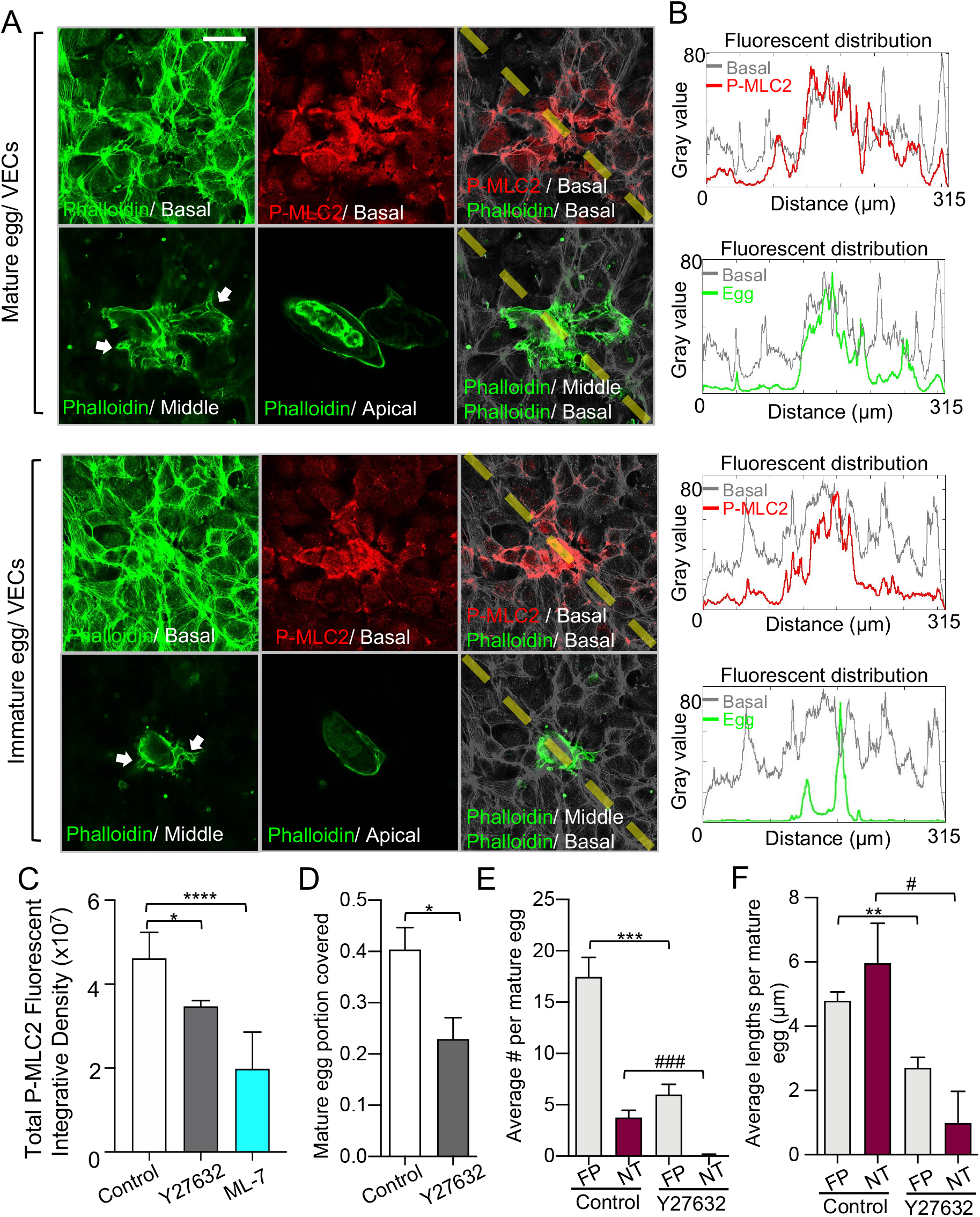
VEC contractility is involved in filopodia formation and encapsulation of eggs. **(A)** Immunostaining of phosphor-MLC2 and F-actin in VECs during encapsulation of live mature eggs at 4 h. The egg location is confirmed by different focal planes of VEC F-actin staining given that the eggshell is covered by VECs. The white arrows in the middle F-actin focal plane show the extended VECs surrounding the egg and indicate the corresponding egg location. **(B)** Fluorescent intensity analysis (the right panel) of phosphor-MLC2 basal (red line)/ F-actin basal (gray line) and F-actin middle (green line)/ F-actin basal (gray line) shows the distributions of phosphor-MLC2 and egg location across the merged images (yellow dashed line). Note how the induction of VEC myosin contractility localizes with the egg location (scale bar: 50 μm). **(C)** Phosphor-MLC2 intensity ratios (mean ± s.e.m.) after 4 h of egg-VEC interaction. VECs were pre-treated with DMSO (white), Y27632 (Rho/RCOCK pathway inhibitor; gray) and ML-7 (myosin light chain kinase inhibitor; blue) for 30 min before incubating with eggs. * and ****, indicate p < 0.01 and p < 0.0001, respectively using a one-way ANOVA with Dunnett’s post-test. Six image fields from three independent experiments were analyzed. **(D)** Quantification of the mature egg portion covered by VECs (mean ± s.e.m.). * indicates p < 0.05 using Student’s *t*-test. **(E)** Quantification of the number (mean ± s.e.m.) of FPs and NT formed per mature egg. *** indicates p < 0.001 and ### indicates p < 0.001 using Student’s *t*-test. **(F)** Quantification of the average lengths of FPs and NTs (mean ± s.e.m.). ** indicates p < 0.01 and # indicates p<0.05 using Student’s *t*-test. For **D** – **F**, 16 and 11 eggs from three independent experiments were used for the control and Y-27632 arms, respectively.

Because the modulation of the cytoskeletal organization by MLC involved the Rho/ROCK pathway (37), we next pre-treated VECs with 20 μM Y-27632 for 30 min and quantified the ability of VECs to encapsulate live mature eggs after 4 h by SEM. Y-27632 decreased by 40% the portion of eggs covered by VECs compared to non-treated VECs (Fig. 4D). Also, Y-27632 decreased the number of filopodia and NTs by 65 and 90%, respectively (Fig. 4E), and the lengths of filopodia and NTs by 40 and 80%, respectively (Figs. 4F). These data indicate the involvement of the Rho/ROCK signaling pathway in the VEC encapsulation process, including the formation of filopodia and NTs.

### Live eggs induce VECs to exert 3D forces to push eggs into the basement substrate

To access the gut lumen and escape from the body, *S. mansoni* eggs must first cross the blood vessel’s basement membrane in the direction perpendicular to the vessel’s wall. The foregoing data regarding encapsulation suggest that mechanical forces may be generated by the VECs in contact with the egg. To investigate this, we used elastic hydrogels (Young’s modulus E = 8 KPa) embedded with 0.2 μm fluorescent microbeads as substrates to culture VECs. These substrates mimic the mechanical characteristics of the native environment of the vessel tunica (38). Measurement of the cell-generated traction forces is accomplished by embedded microspheres and then tracking their displacements. 3D traction force microscopy (3D-TFM) is used to calculate the 3-D traction forces from microsphere displacements (31, 39, 40). After a 24 h of interaction between a live mature egg and VECs, we imaged the displacement of the fluorescent microbeads in the hydrogel substrates at different focal planes (Fig. 5A). The surface microbeads were pushed down to the middle focal plane as the egg volume deformed the subendothelial elastic hydrogel. The XZ projection showed the bending surface of the subendothelial layer over the 24 h interaction (Fig. 5A).

**Figure 5.**
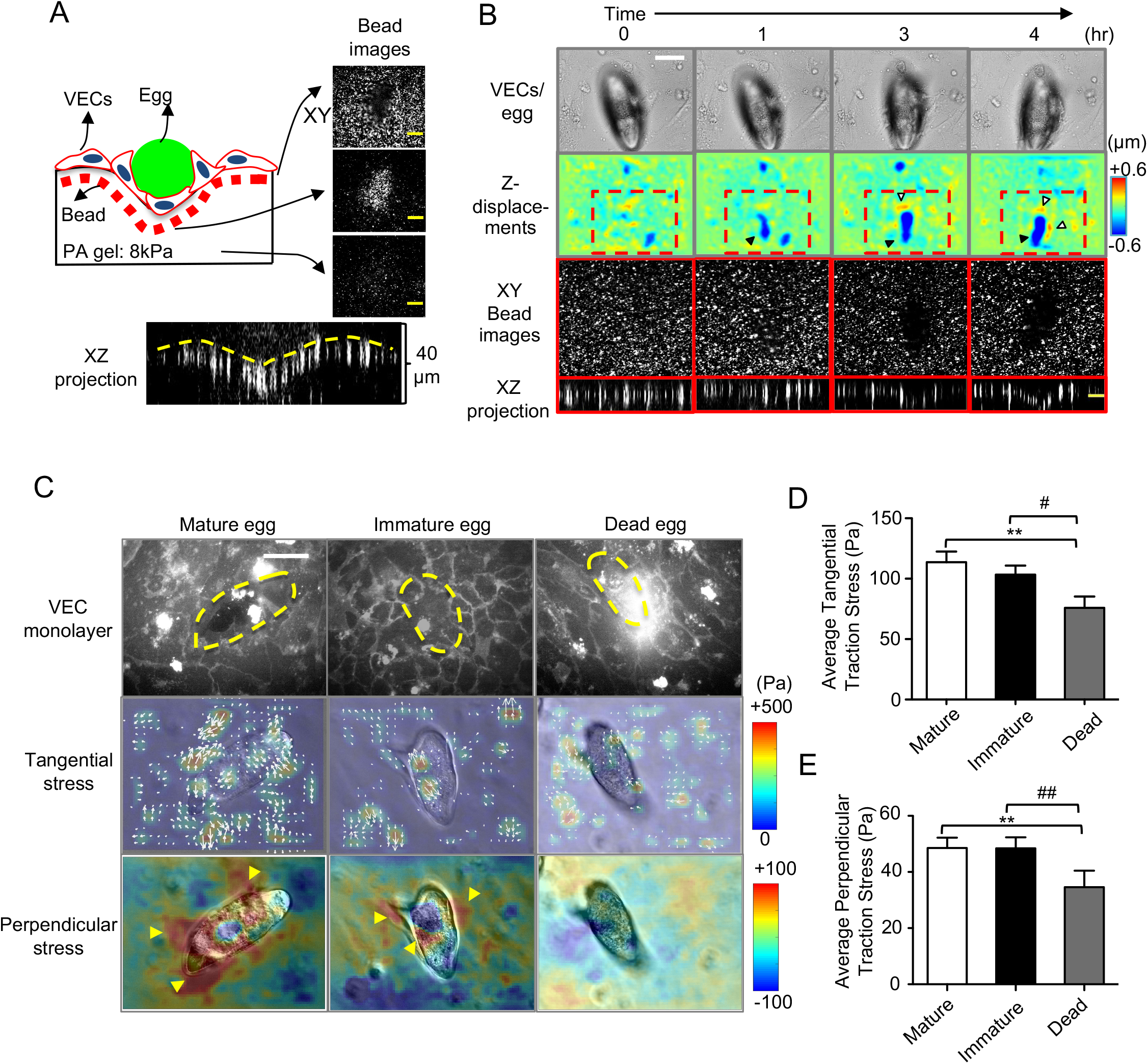
3D traction forces exerted by VECs during encapsulation of *S. mansoni* eggs. **(A)** Schematic of the 3D confocal microscopy setup to measure the 3D deformation of VECs during encapsulation. Fluorescent beads (red) were embedded in the elastic FN-coated elastic hydrogel substrates on which the VECs and eggs (green) were co-cultured. Orthogonal (xy and xz) views of the beads’ fluorescent channel demonstrate the 3D substrate deformation underneath the eggs (scale bar: 10 μm). **(B)** Temporal evolution of substrate deformation during encapsulation of a live mature egg. The top row shows bright-field images of the egg and VECs (scale bar: 50 μm). The second row shows maps of the z-displacement (perpendicular to the substrate) in the same field of view. Downward pushing patterns (negative z-displacement) are indicated by black arrowheads whereas upward pulling patterns (positive z-displacement) are indicated by white arrowheads. The third and fourth rows, respectively, show xy and xz enlarged images of beads in those areas with the greatest deformation (scale bar: 10 μm) as delimited by the red dashed-line box in the second row. **(C)** 3D traction stresses exerted by VECs interacting with live mature, live immature and dead eggs after 4 h. The top row shows VEC cell membrane staining (scale bar: 50 μm). The yellow dashed line shows the egg location. The second row shows the in-plane tangential (xy) stress magnitude (represented by the color map) and direction (white arrows). The third row shows the perpendicular (xz) stress magnitude. Positive/negative values indicate upward pulling and downward pushing, respectively. Note the downward pushing forces (blue color) localized at the egg anchoring site which is surrounded by upward pulling forces (red color, also indicated by yellow arrowheads). **(D-E)** Quantification of tangential **(D)** and perpendicular **(E)** traction stress magnitudes. Data are presented as the mean ± s.e.m. ** indicates p < 0.01, and # and ## indicate p < 0.05 and p < 0.01, respectively, using one-way ANOVA with Turkey’s post-test. For mature, immature and dead eggs, 26, 20 and 12 eggs were used, respectively, across three independent experiments.

Next, we performed a time course experiment to investigate the dynamic changes of the VEC monolayer and subendothelial substrate in response to a live mature egg. After a 1 h interaction between egg and VEC, the substrate deformation had increased by over 2-fold in the z-direction perpendicular to the monolayer compared to zero-time point (Fig. 5B). After 3 and 4 h, the net z-deformations of the substrate by the egg had increased by up to 4-fold (SI Fig. 4 and Movie 3). Such downward deformations measured under the *S. mansoni* egg suggest that VECs actively generate 3D forces to push the egg into the tissue. To study whether egg vitality affects the force generated by VECs, we used 3D-TFM to quantify the forces exerted by VECs in contact with live mature and immature eggs, and dead eggs for 4 h (Fig. 5C). The 3D distributions of the traction forces data showed that VEC tangential forces in the x-y direction had an irregular distribution of hotspots where the VECs were in direct contact with the egg (the second row of Fig. 5C). Such force patterns have also been observed underneath endothelial monolayers when VECs interact with functionalized beads (30, 31). In contrast, the perpendicular forces in the z-direction had a more defined pattern with a downward pushing zone at the egg anchoring point and an upward pulling zone surrounding it (yellow arrowheads in the third row of Fig. 5C). At 4 h, both the tangential and perpendicular forces were significantly stronger for mature and immature eggs relative to dead eggs, indicating that egg vitality is an important contributor to the process (Figs. 5D–E).

### Increases in VEC monolayer tension during encapsulating cause VEC junctions to rupture

We examined whether the increased 3D deformations caused by the contact between VECs and parasite eggs resulted in endothelial barrier disruption. Specifically, we performed 3D confocal microscopy of immunostained VE-cadherin, the major molecule that regulates endothelial barrier function (41), during encapsulation (Fig. 6A). The VEC monolayers co-cultured with eggs for 4 h had disrupted VE-cadherin junctional connections that co-localized with sites of downward pushing and egg-hydrogel contacts. Moreover, VEC permeability was significantly increased in the presence of live mature eggs compared to either dead eggs or eggshells (SI Fig. 5).

**Figure 6.**
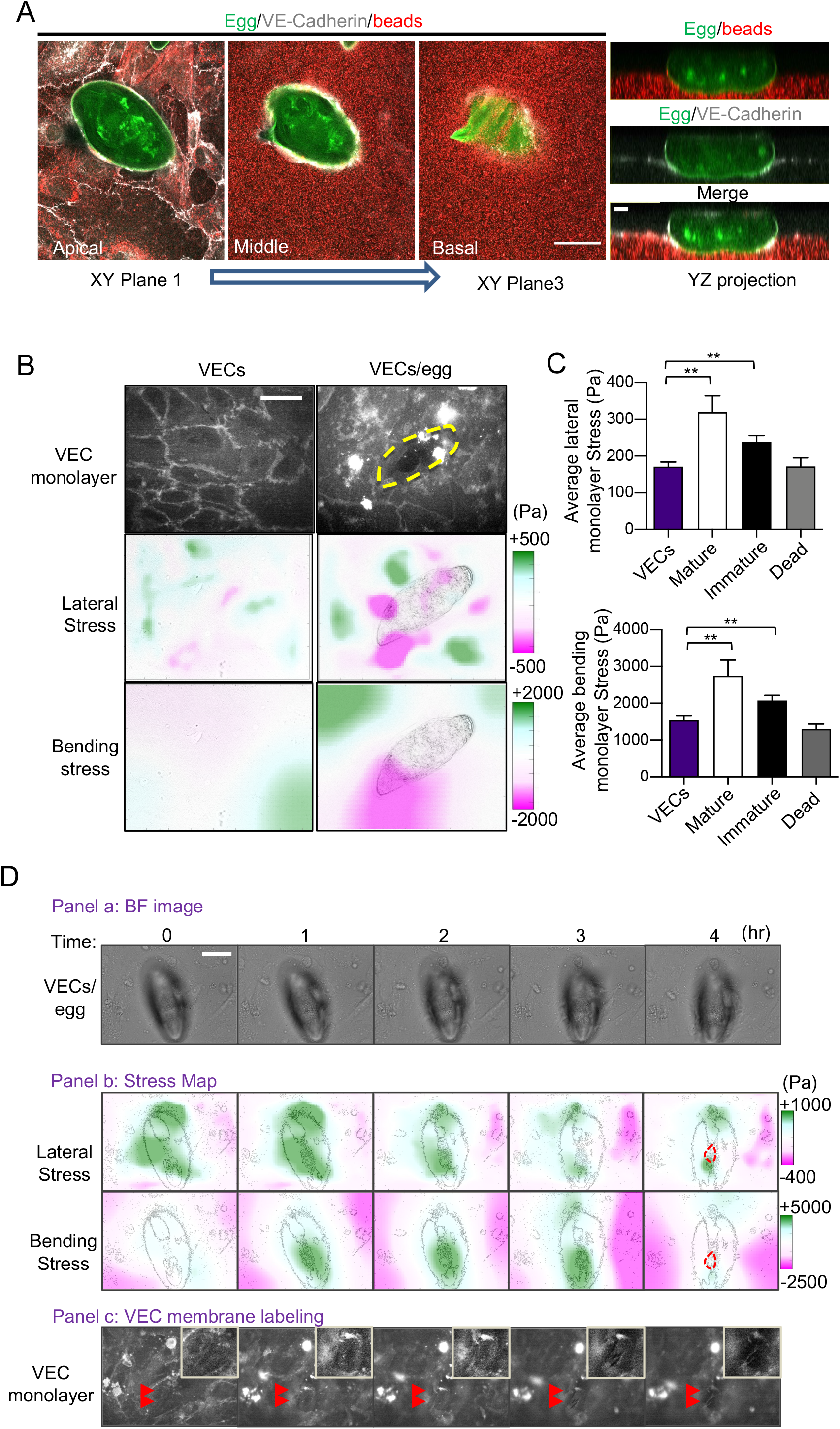
Increased monolayer tension during the encapsulation of *S. mansoni* ruptures VEC junctions. **(A)** Confocal images of a live mature egg being encapsulated by VECs after 4 h on a PA gel seeded with fluorescent beads. The three left-hand-side panels show xy images of VE-Cadherin staining (white), the egg (green) and beads within the gel (red), taken at the monolayer z-plane and then with increasing depth into the substrate (scale bar: 50 μm). The right-hand-side panel shows the corresponding yz projection (scale bar: 10 μm). **(B)** 3D stress patterns exerted by VEC monolayers in the presence and absence of a live mature egg after 4 h. The top row shows VEC junctions (scale bar: 50 μm). The yellow dash line outlines the egg location. The second and third rows represent lateral and bending monolayer tensions, respectively. **(C)** Quantification of the lateral (upper graph) and bending monolayer tension magnitudes (mean ± s.e.m.). Control VECs without eggs (n=10), and VECs with 25, 20 and 12 live mature and immature, and dead eggs from three independent experiments, respectively, were analyzed. ** indicates p < 0.01 using Welch’s *t*-test. **(D)** Temporal evolution of the VEC monolayer during the encapsulation of a live mature egg. Panel a (bright field images): the first row shows images of the egg and VECs (scale bar: 50 μm). The second row shows VEC junctions in the same field of view. Red arrowheads indicate an area where a junctional gap between the VECs has opened up underneath the egg (top right corner inserts show enlarged images of the region of interest). Panel b (stress maps): the first and second rows, respectively, show lateral and bending monolayer stresses superimposed on the egg’s silhouette. The red dashed contour at t = 4 h indicates the VEC gap location.

To gain insight into the mechanism for endothelial disruption, we employed 3D monolayer stress microscopy (3D-MSM) (42) to quantify the changes in VEC monolayer tensions elicited by live *S. mansoni* eggs. The transmission of VEC contractile forces at cell-cell contacts raises the intracellular tension of the VEC monolayer by laterally stretching and bending the monolayer, and this tension can modulate the monolayer barrier function (43, 44). Our measurements revealed that VEC monolayers interacting with live mature and immature eggs generated greater lateral and bending tensions compared to monolayers without eggs (Fig. 6B). These differences in tension were significant in the lateral (Fig. 6C, upper panel; monolayer *vs.* mature, p = 0.0032; monolayer *vs.* immature, p = 0.0036) and vertical (bending) directions (Fig. 6C, lower panel; monolayer *vs.* mature, p = 0.0083; monolayer *vs.* immature, p = 0.0015): no statistical differences were measured between dead eggs and the monolayers alone.

Finally, over a 4 h period of VECs interacting with a mature egg, we measured the monolayer tension during which the VEC junctions underneath the egg became dissociated to eventually form a gap (Fig. 6D panels a and c). Lateral tension (Fig. 6D, panel b, 1st row) was initially strong (t = 0) and remained elevated through the first two hours of interaction but decreased to more fluctuating patterns as the junctions began to dissociate at 3 h. On the other hand, the bending tension (Fig. 6D, panel b, 2nd row) was initially weak (t = 0) but increased markedly over the next three hours as VECs encapsulated the egg and pushed it downwards. Eventually, the occurrence of a monolayer gap at the 4 h time point disrupted both the lateral and bending monolayer tension by creating a stress-free boundary along the gap perimeter. Such breaks in the integrity of the junctions were noted in approximately 70% of the live egg-VEC interactions observed. These data imply that the VEC monolayer actively alters its biomechanical state when in contact with live S. *mansoni* eggs and this, in turn, disrupts the integrity of VEC junctions and facilitates passage of the egg. The time course of the data suggests that the increase in bending of the monolayer due to the egg being pushed into the basal membrane precedes loosening of the VEC junctions.

## Discussion

Migration through tissues by *S. mansoni* eggs is an essential step in the parasite’s life cycle. Contact between the parasite’s eggs and VECs triggers intravascular host-immune responses to induce VEC inflammation, proliferation and migration (6–8, 45, 46). However, the biomechanical mechanisms regulating egg-VEC interactions are unknown. We used quantitative microscopy to show that S. *mansoni* eggs stimulate VECs to form membrane protrusions that facilitate the encapsulation of eggs. When VECs came into contact with eggs, they exerted 3D mechanical forces mediated by the activation of Rho/ROCK signaling that push the egg towards the basement membrane. The increased VEC contractility also caused monolayer tension to nearly double and fluctuate, thus, destabilizing cell-cell junctions and facilitating egg extravasation. Importantly, we demonstrate that the mechanisms described are enhanced by egg vitality, whereby dead eggs are far less stimulatory. Overall, our data describe a mechanism by which *S. mansoni* eggs hijack the contractile machinery of the host’s endothelium to propel their motion toward the extravascular space.

The eggshell of *S. mansoni* is composed of chitin and has a dense distribution of surface projections, commonly referred to as microspines (see (16, 20, 21) and references therein). We studied the interactions between the egg microspines and VEC membrane protrusions at the nanometer scale using SEM. Filopodia are thin, actin-rich bundled fibers that protrude from cell membranes and serve several physiological roles, such as probing the environment and facilitating cell motility (47). Apart from filopodia, recent studies have identified another type of cell membrane protrusion, named intercellular NTs or tunneling NTs, which do not attach to the substrate but establish connections between cells and are thought to mediate long-range intercellular communication (35). These NTs can transfer cytoplasmic material and pathogens such as HIV from one cell to another (36, 48). In VECs, the formation of intercellular NTs can be stimulated by oxidative stress, apoptosis, and inflammation (49, 50). Interestingly, we observed that VECs generated both filopodia and NTs when they were in direct contact with S. *mansoni* eggs. Filopodia appeared at the front of the “leader” VECs to probe the egg surface, whereas intercellular NTs were found at the trailing edge of these leader cells to interconnect with the basal endothelium. Filopodia extended from the VEC cytoskeleton and firmly adhered to the eggshell, and the number of these thin frontal protrusions positively correlated with the eggshell area covered by the VECs.

Collective cell migration relies on the long-range coordination of cell polarity between leader and follower cells (51). Our data suggest that filopodia and NTs contribute to regulating collective cell polarity during egg encapsulation. We determined that the leading filopodia on VECs exhibited a ‘seeking’ or ‘exploration’ phenotype, advancing to adhere to a new egg before completely covering those that were already engaged by VECs (Fig. 2B). Adherens junctions have been implicated as major contributors to collective polarity by transmitting cellular forces across flat monolayers (29). In contrast to adherens junctions, NTs mediate long-range collective communication over distances of up to 100 μm in leukocytes (52). We observed endothelial cell NTs as long as 20 μm in our experiments mediating long-range cell-cell connections during VEC migration over eggs. Thus, we propose that NTs could reinforce the collective VEC behavior during encapsulation in a manner similar to their functionality in the embryonic cell sheet-like migration model during which leader cells at the migratory front extend intercellular NT bridges to help pull the cells behind while maintaining contact (53).

Immature eggs are released from the female worm onto the mesenteric veins to interact with VECs. During their migration through the extravascular tissues, the developing embryo is separated from the eggshell by the Reynold’s layer (12). Microscopic pores are present within the rigid eggshells (54, 55) through which ESPs from the embryo are secreted (12). The analysis of ESPs from immature eggs is less detailed; however, it is known that freshly deposited immature eggs induce a faster VEC encapsulation process without inducing granuloma formation (6, 56). In contrast, mature egg ESPs are believed to activate VEC proliferation, migration and angiogenesis, and recruit leukocytes to initiate the granulomatous response which facilitates egg migration (6–8, 25, 57). For example, internalization of the major mature egg ESP, Interleukin (IL)-4-inducing principle, by VECs has been associated with egg-induced endothelial proliferation, angiogenesis and blood vessel leakiness (58, 59). These findings including together with our data suggest that ESPs from immature and mature eggs interact with VECs at different egg extravasation stages. We hypothesize that immature egg ESPs might induce a fast VEC encapsulation response in the first 24 h. Subsequently, the encapsulated eggs lodge between endothelial and basement membrane layers for 4-5 days to mature, remaining in contact with the VECs to initiate the early granulomatous inflammation. This is consistent with the knowledge that eggs must be fully mature to migrate from the blood vessel into the gut (56, 60), a process that can take approximately one to four weeks (26).

The present data indicate that live eggs increase VEC migratory activity. VECs encapsulated both mature and immature live eggs at similar rates covering ~40% of the eggshell surface within the first 4 h of interaction, whereas the encapsulation of dead eggs was significantly less prominent (~20%) over the same period. Also, the VECs interacting with dead eggs had ~2.5-fold fewer filopodia and ~10-fold fewer intercellular NTs than those interacting with live eggs. In addition to vitality, the egg’s developmental state influenced the dynamics of filopodia and NT formation. Thus, the filopodia on dead and live mature eggs had similar lengths (~4-5 μm), whereas those on live immature eggs were twice as long. We also found that VECs migrated slightly faster over live immature eggs compared to mature eggs (13-18 μm vs. 8-13 μm in the z-direction between 0.5 and 1 h). We speculate that the longer VEC filopodia on immature eggs contribute to faster VEC encapsulation and could be mediated by both biochemical and biophysical mechanisms, such as a different spectrum of ESPs secreted from immature eggs, metabolic enzymes secreted from adult worms that attached to the eggshell (4, 61), and/or the eggshell surface topography (SI Fig. 3).

During cell migration, actomyosin-based contractility integrates the filopodial F-actin bundles into protruding lamellipodia at the cell front (47). The Rho/ROCK pathway is a key regulator of cell migration that coordinates cell polarity and contractility (37). Specifically, triggering of the Rho/ROCK pathway activates myosin II, which mediates cytoskeletal remodeling and modulates intracellular tension to control the dynamics of cell adhesion (62). In this context, our experiments showed that MLC phosphorylation was enhanced in those VECs that were in direct contact with *S. mansoni* eggs, implying that myosin II contractility had been activated. Moreover, small molecule inhibition of the Rho kinase activity in VECs decreased their ability to encapsulate eggs, as well as the length and number of filopodia, and intercellular NTs. These results highlight the importance of the VEC Rho/ROCK pathway in driving VEC motility during the early stages of egg extravasation. Furthermore, our data suggested that *S. mansoni* eggs modulate the contractile machinery of the surrounding vascular cells, which motivated us to investigate whether the resulting forces generated by the VECs are sufficient to physically drive the eggs across the endothelium.

S. *mansoni* eggs cannot move on their own and must rely on external forces to transit from the vasculature toward the intestinal lumen. Given the egg’s voluminous size and rigidity (dimensions ~100 × 30 × 30 μm and a Young’s modulus ~10 MPa), this extravasation process likely causes large deformations in the egg’s surroundings and requires significant forces. The origin of these forces has been the subject of speculation in the past, leading to two main theories. One theory suggests that the eggs recruit endothelial cells to push them into the basal membrane and promote proteolysis of the extracellular matrix (6, 26). An alternate theory proposes that muscular contractions of the female worm force the eggs into the vessel wall (27). To shed light on these questions in relation to eggs, we used 3D-TFM to measure the external forces involved during the interaction of VECs with eggs *in vitro*. Using pliant elastic hydrogels embedded with 0.2 μm fluorescent microspheres as cell substrates, we mimicked the mechanical properties of the egg’s physiological environment (63). Significant substrate indentations and bending of the VEC monolayer occurred under the eggs during the first hour of encapsulation, and these increased in magnitude over time. The tangential (in-plane) traction stresses exerted by the VECs increased ~1.6-fold under mature and immature eggs compared to dead eggs, showing a disordered vector pattern. In addition, perpendicular stresses also increased ~1.6-fold and appeared under the eggs, displaying a peripheral ring of upward pulling traction surrounding a small area of intense downward pushing (SI Fig. 6). This pattern of 3D traction stress is consistent with that observed at cell-cell junctions during the engulfment by VECs of inert particles coated with the anti-intercellular adhesion molecule (ICAM) (31). Given that VEC ICAM expression is strongly upregulated upon exposure to egg ESPs (25), we posit that eggs activate VECs to generate forces strong enough to push the eggs across the endothelium. Finally, the large lateral spine of the S. *mansoni* egg has been speculated to play a role in egg extravasation (64). Accordingly, we analyzed the substrate indentation sites and the egg spine positions (SI Fig. 7) and found that they did not co-localize. Thus, it seems that the egg spine does not contribute to the initial stages of egg extravasation.

The ability of the *S. mansoni* egg to promote 3D forces from VECs depends, to a quantitative degree, on the vitality of the *S. mansoni* egg. Specifically, we found that VECs in contact with live eggs generated traction stresses that were at least twice as strong as those generated by VECs in contact with dead eggs. The VEC monolayers in contact with live eggs also had a larger intracellular tension than those in contact with dead eggs. Under increased tension, VEC monolayers have loosened adherens junctions and increased migration, leading to transient endothelial gaps (30, 31). Indeed, we found that live eggs often caused the breakdown of VEC junctions with an increase in endothelial permeability. Thus, we postulate that the modulation of VEC contractility and monolayer tension are orchestrated by secretions from the live egg, which not only force the egg into the vessel wall but also shift the VEC monolayer to a state of decreased barrier function. Together, our findings demonstrate how the biomechanical interactions between.*S. mansoni* eggs and VECs are critical for successful egg extravasation.

Successful passage of the *S. mansoni* egg from the endovascular space to the lumen of the gut is essential for continuation of the parasite’s lifecycle. Mature eggs are the only form that the human sheds into the environment, highlighting the importance of egg vitality in the extravasation process (56). Although it is well-established that schistosome eggs are biologically active and release factors that modulate the immune response during tissue migration (cited above), the biomechanical processes at play in the first and crucial steps of egg encapsulation and extravasation, have been less studied. Our investigations comparing live mature and immature, and dead *S. mansoni* eggs, have discovered how egg vitality contributes to activating VECs to encapsulate and exert 3D mechanical forces on eggs that eventually disrupt VEC junctions to transport eggs away from the endothelial surface into the underlying tissues. Future work will examine specific components of the ESPs for their possible contributions to the biomechanical processes described and measured here, including possible differences between schistosome species.

## Materials and Methods

### Reagents

The VE-Cadherin antibody used for immunostaining was purchased from Santa Cruz Biotechnologies (sc-9989). The antibody against phosphor-MLC2 serine 19 was purchased from Cell Signaling Technology (3671). Alexa Fluor-488-, Alexa Fluor-594-Phalloidin and Cell Mask deep red plasma membrane stain used for immunostaining and/or live cell labeling were purchased from Invitrogen (A12379, A12381 and c10046). The carboxylate-modified red (580/605) microspheres used for traction force microscopy were purchased from Invitrogen (F8809). Y27632 was purchased from Abcam (ab120129) and ML-7 was purchased from Sigma (475880).

### Ethics Statement

Male Golden Syrian hamsters infected with the Naval Medical Research Institute (NMRI, Puerto Rican) isolate of *S. mansoni* were obtained from the Biomedical Research Institute (BRI, Rockville, MD) (65) and maintained in accordance with protocols approved by the Institutional Animal Care and Use Committee (IACUC) at the University of California San Diego.

### Isolation of *S. mansoni* eggs

Eggs were isolated from the livers of hamsters six weeks after infection with 600 *S. mansoni* cercariae (66). Specifically, livers were finely minced with a sterile razor blade and digested overnight at 37°C in 40 mL 1×PBS containing 2% penicillin/streptomycin/amphotericin B (1×PBS −2% PSF) and 5 mL 0.5% clostridial collagenase solution (0.025 g of collagenase (Sigma, C0130) in 5 mL of dH_2_0 for immediate use). All subsequent steps proceeded at room temperature. The digested livers were centrifuged at 400 × *g* for 5 min and the supernatant was decanted. The resulting pellet was resuspended in 50 mL 1×PBS-2% PSF, centrifuged under the same conditions and the supernatant decanted: this step was repeated 3-5 times until the supernatant was clear. During this process, eggs settle to the bottom of the conical tube. Eggs were layered gently onto the first Percoll gradient (8 mL sterile Percoll with 32 mL 0.25 M sucrose) and centrifuged at 800 × *g* for 10 min to separate any remaining liver tissue debris. The supernatant was discarded with a serological pipette. Eggs were resuspended in 3 mL 1×PBS −2% PSF and gently applied to the second Percoll gradient (2.5 mL Percoll with 7.5 mL 0.25 M sucrose). Immature eggs were removed at the interface and mature eggs were collected from the bottom of the column. Immature eggs were further separated using a third Percoll gradient (6 mL of Percoll, 0.6 mL 9% saline and 3.4 mL M199 culture medium) and centrifugation for 15 min at 250 × *g* (12). The mature and immature egg fractions were washed 3-5 times with M199 medium, centrifuged at 310 × *g* for 3 min to remove any remaining Percoll and tissue debris. Mature egg fractions typically contain less than 5% immature eggs, whereas immature egg fractions contain 5-10% mature eggs, as observed microscopically. Hatching of mature eggs in water was usually ~80% after 40 min under a bright light, thus confirming their viability. To generate dead eggs, mature eggs were treated with 1% sodium azide (Sigma S2002) for 24 h in M199 and then washed six times in the same medium. All experiments were conducted within five days of preparing eggs.

### Cells and co-culture

Human umbilical VECs (Cell Application 200-05n) were cultured on fibronectin (Sigma 10838039001)-coated glass slides or polyacrylamide (PA) hydrogels for 48 h in M199 supplemented with 10% endothelial cell growth medium (Cell Application), 10% FBS (Lonza), 1% sodium pyruvate, 1% L-glutamine and 1% penicillin-streptomycin (Gibco) until they formed a confluent monolayer. Eggs were added on top of the VECs at a density of 4 eggs /mm^2^ and co-incubated for up to 24 h.

### Immunofluorescence and confocal microscopy

Cells were cultured on fibronectin-coated substrates (50 μg/mL) and treated with eggs or reagents at the indicated time points. After treatment, cells were fixed in 4% paraformaldehyde for 10 min. Cells were then permeabilized in 1×PBS supplemented with 0.1% Triton X-100 for 10 min followed by a blocking step in 1×PBS supplemented with 5% BSA (Sigma A7906). Cells were incubated with primary and secondary antibodies and, after each step, washed with 1×PBS. DAPI (Sigma 10236276001, 1μg/mL) was used as a counterstain. Images were taken with an epi-fluorescence microscope (Olympus IX70) or a confocal microscope (Zeiss 880 Airyscan). For 3D-TFM, z-stack image acquisition was performed using a spinning disc confocal microscope (Olympus IX81), a 40X NA 1.35 oil immersion objective lens, a cooled CCD (Hamamatsu) camera and the Metamorph software version 7.8.10 (Molecular Devices). For the 3D image analysis, sequences of taken z-stack images were taken and analyzed using the Volocity software version 6.3 (Quorum Technologies Inc.) which rendered the optical sections as 3D models, thus enabling the analysis of the interactions between parasite eggs and VECs.

### Scanning Electron Microscopy (SEM)

Samples were fixed with 2% glutaraldehyde (Electron Microscopy Sciences) and 4% paraformaldehyde, pH 7.4, for 24 h. Samples were critical point dried in increasing concentrations of high-grade ethanol using an Autosamdri 815 critical point dryer and then sputter coated with iridium using an Emitech K575X. Imaging was done with a QUANTA FEG 250 ESEM (Field Electron and Ion Company) or Sigma 500 SEM (Zeiss). For each image, the average length and number of filopodia and intercellular NTs was measured and counted using the FIJI imaging analysis software (67).

### Polyacrylamide (PA) gel preparation and characterization

On 35-mm glass-bottom dishes (World Precision Instrument FD35-100), we fabricated 12 mm diameter and 40 μm thick PA gels in 1×PBS by first mixing 5% acrylamide and 0.3% bis-acrylamide (Young’s modulus E = 8.7 KPa) and then adding a 1/100 total volume of 10% APS and a 1/1000 total volume of TEMED (Sigma; to initiate gel polymerization). To improve the signal to noise ratio of the z-stack images and the displacement field calculation, the PBS used in the fabrication of the gels contained 0.03% carboxylate-modified red microspheres (0.2 μm diameter; Thermo Fisher Scientific). After placing a coverslip on top, the side was inverted and the gel allowed to polymerize for 30 min, during which time the microbeads migrated to the surface of the gel. The distribution of the microbeads at the surface of the gels was verified by the 3D confocal microscopy. To encourage cell attachment to the PA gels, we crosslinked the extracellular matrix protein, fibronectin, to the gel surface by using UV activated Sulfo-SANPAH (0.15 mg/mL; Thermo Scientific 22589). The gels were incubated overnight at 4 °C.

### Three-Dimensional Traction Force Microscopy (3D-TFM) and Monolayer Stress Microscopy (MSM)

The 3D deformation of the PA gel’s surface, in which the fluorescent microspheres were localized, was measured with a confocal microscope (Olympus IX81). The 3D deformation field was formulated as ***u***(*x*, *y*, *z* = *h*), where the deformation vector ***u*** depends on the coordinates of *x*, *y*, *z* and for a single horizontal plane measurement, *z* = *h*. Time-lapse sequences of fluorescence z-stacks consisting of 40 planes separated by 0.4 μm were acquired at 1 h intervals. The 3D deformation was determined by cross-correlating each instantaneous z-stack with an undeformed reference z-stack taken after detaching the cells by trypsin treatment. After imaging, attached cells were trypsinized and their removal relaxed the elastic substrates back to the undeformed state., that served as reference for the correlations. To balance the spatial resolution and signal-to-noise ratio, the *z-*stacks were divided into 3D interrogation boxes of 32 × 32 × 12 pixels in the *x*, *y*, and *z* directions, respectively. These settings provided a Nyquist spatial resolution of 2 μm in three spatial directions. The elasticity equation of equilibrium was solved for a linear, homogeneous and isotropic body with a Poisson’s ratio σ=0.45 (39, 40, 68) to determine the 3D deformation everywhere inside the substrate, ***u***(*x*, *y*, *z*), from ***u***(*x*, *y*, *h*). We then applied Hooke’s law to calculate the six independent components of the stress tensor everywhere inside the substrate. In particular, we computed the 3D traction stress vector at the surface of the substrate in contact with the cells, [*τ*_*xy*_(*x*, *y*, *h*), *τ*_*yz*_(*x*, *y*, *h*), *τ*_*zz*_(*x*, *y*, *h*)]. We used 3D MSM to infer the intracellular tension caused by lateral deformation and bending of cell monolayers (42). The calculation is carried out by imposing equilibrium of forces and moments in the monolayer subject to external loads given by the 3D traction stresses.

### Statistical analysis

For comparisons between two groups, statistical analyses were performed by two-tailed unpaired Student’s *t-*test and Welch’s *t*-test. Comparison of multiple groups was made by one-way ANOVA, and statistical significance among multiple groups was determined by Tukey’s post-test or Dunnett’s post-test (for pair-wise comparisons of means). All results are presented as means ± s.e.m. from three independent experiments.

## Acknowledgements

We thank James H. McKerrow of the CDIPD, Skaggs School of Pharmacy and Pharmaceutical Sciences, University of California San Diego for advice.

## References

1. WHO Expert Committee. Prevention and control of schistosomiasis and soil-transmitted helminthiasis.: World Health Organ Tech Rep Ser.; 2002. 57 p.

2. Loker ES, Hofkin, BV, Mkoji, GM, Mungai, B, Kihara, J, Koech, DK. Distributions of freshwater snails in southern Kenya with implications for the biological control of schistosomiasis and other snail-mediated parasites. J Med Appl Malacol. 1993;5:20.

3. Costain AH, MacDonald AS, Smits HH. Schistosome Egg Migration: Mechanisms, Pathogenesis and Host Immune Responses. Front Immunol. 2018;9:3042.

4. Schwartz C, Fallon PG. Schistosoma “Eggs-Iting” the Host: Granuloma Formation and Egg Excretion. Front Immunol. 2018;9:2492.

5. Boros DL, Warren KS. Delayed hypersensitivity-type granuloma formation and dermal reaction induced and elicited by a soluble factor isolated from Schistosoma mansoni eggs. J Exp Med. 1970;132(3):488–507.

6. File S. Interaction of schistosome eggs with vascular endothelium. J Parasitol. 1995;81(2):234–8.

7. Lejoly-Boisseau H, Appriou M, Seigneur M, Pruvost A, Tribouley-Duret J, Tribouley J. Schistosoma mansoni: in vitro adhesion of parasite eggs to the vascular endothelium. Subsequent inhibition by a monoclonal antibody directed to a carbohydrate epitope. Exp Parasitol. 1999;91(1):20–9.

8. Loeffler DA, Lundy SK, Singh KP, Gerard HC, Hudson AP, Boros DL. Soluble egg antigens from Schistosoma mansoni induce angiogenesis-related processes by up-regulating vascular endothelial growth factor in human endothelial cells. J Infect Dis. 2002;185(11):1650–6.

9. Homeida M, Ahmed S, Dafalla A, Suliman S, Eltom I, Nash T, et al. Morbidity associated with Schistosoma mansoni infection as determined by ultrasound: a study in Gezira, Sudan. Am J Trop Med Hyg. 1988;39(2):196–201.

10. Dessein AJ, Hillaire D, Elwali NE, Marquet S, Mohamed-Ali Q, Mirghani A, et al. Severe hepatic fibrosis in Schistosoma mansoni infection is controlled by a major locus that is closely linked to the interferon-gamma receptor gene. Am J Hum Genet. 1999;65(3):709–21.

11. Cheever AW. A quantitative post-mortem study of Schistosomiasis mansoni in man. Am J Trop Med Hyg. 1968;17(1):38–64.

12. Ashton PD, Harrop R, Shah B, Wilson RA. The schistosome egg: development and secretions. Parasitology. 2001;122(Pt 3):329–38.

13. Vogel H. Über Entwicklung, Lebensdauer und Tod der Eier vom Bilharzia japonica im Wirtsgewebe. Dtsch Tropenmed Ztsch. 1942;46:35.

14. Prata A. Biópsia retal na esquistossomose mansoni — bases e aplicações no diagnóstico e tratamento. Serviço Nacional de Educação Sanitária-Ministério da Saúde, Rio de Janeiro. 1957.

15. Jurberg AD, Goncalves T, Costa TA, de Mattos AC, Pascarelli BM, de Manso PP, et al. The embryonic development of Schistosoma mansoni eggs: proposal for a new staging system. Dev Genes Evol. 2009;219(5):219–34.

16. Basch PF. Schistosomes: Development, Reproduction and Host Relations. Oxford University Press. 1991.

17. Eissa MM, El Bardicy S, Tadros M. Bioactivity of miltefosine against aquatic stages of Schistosoma mansoni, Schistosoma haematobium and their snail hosts, supported by scanning electron microscopy. Parasit Vectors. 2011;4:73.

18. Inatomi S. Submicrospic structure of the eggshell of helminths Journal of Okayama Medical Association. 1962;74:51.

19. Stenger RJ, Warren KS, Johnson EA. An ultrastructural study of hepatic granulomas and schistosome egg shells in murine hepatosplenic schistosomiasis mansoni. Exp Mol Pathol. 1967;7(1):116–32.

20. Ford JW, Blankespoor HD. Scanning electron microscopy of the eggs of three human schistosomes. Int J Parasitol. 1979;9(2):141–5.

21. Karl S, Gutierrez L, Lucyk-Maurer R, Kerr R, Candido RR, Toh SQ, et al. The iron distribution and magnetic properties of schistosome eggshells: implications for improved diagnostics. PLoS Negl Trop Dis. 2013;7(5):e2219.

22. Hoeppli R. Parasitic and free-living nematodes found on the Island of Amoy. Report M R A C. 1932.

23. Cass CL, Johnson JR, Califf LL, Xu T, Hernandez HJ, Stadecker MJ, et al. Proteomic analysis of Schistosoma mansoni egg secretions. Mol Biochem Parasitol. 2007;155(2):84–93.

24. Mathieson W, Wilson RA. A comparative proteomic study of the undeveloped and developed Schistosoma mansoni egg and its contents: the miracidium, hatch fluid and secretions. Int J Parasitol. 2010;40(5):617–28.

25. Ritter DM, McKerrow JH. Intercellular adhesion molecule 1 is the major adhesion molecule expressed during schistosome granuloma formation. Infect Immun. 1996;64(11):4706–13.

26. deWalick S, Tielens AG, van Hellemond JJ. Schistosoma mansoni: the egg, biosynthesis of the shell and interaction with the host. Exp Parasitol. 2012;132(1):7–13.

27. Linder E. The Schistosome Egg in Transit. Annals of Clinical Pathology. 2017;5(3).

28. Zhang S, Skinner D, Joshi P, Criado-Hidalgo E, Yeh YT, Lasheras JC, et al. Quantifying the mechanics of locomotion of the schistosome pathogen with respect to changes in its physical environment. J R Soc Interface. 2019;16(150):20180675.

29. Charras G, Yap AS. Tensile Forces and Mechanotransduction at Cell-Cell Junctions. Curr Biol. 2018;28(8):R445–R57.

30. Hur SS, del Alamo JC, Park JS, Li YS, Nguyen HA, Teng D, et al. Roles of cell confluency and fluid shear in 3-dimensional intracellular forces in endothelial cells. Proc Natl Acad Sci U S A. 2012;109(28):11110–5.

31. Yeh YT, Serrano R, Francois J, Chiu JJ, Li YJ, Del Alamo JC, et al. Three-dimensional forces exerted by leukocytes and vascular endothelial cells dynamically facilitate diapedesis. Proc Natl Acad Sci U S A. 2018;115(1):133–8.

32. Lamason RL, Bastounis E, Kafai NM, Serrano R, Del Alamo JC, Theriot JA, et al. Rickettsia Sca4 Reduces Vinculin-Mediated Intercellular Tension to Promote Spread. Cell. 2016;167(3):670–83 e10.

33. Bastounis EE, Alcade FS, Radhakrishnan P, Engstrom P, Benito MJG, Oswald MS, et al. Mechanical competition triggered by innate immune signaling drives the collective extrusion of bacterially-infected epithelial cells. bioRxiv. 2020.

34. Rustom A, Saffrich R, Markovic I, Walther P, Gerdes HH. Nanotubular highways for intercellular organelle transport. Science. 2004;303(5660):1007–10.

35. Gerdes HH, Carvalho RN. Intercellular transfer mediated by tunneling nanotubes. Curr Opin Cell Biol. 2008;20(4):470–5.

36. Gerdes HH, Bukoreshtliev NV, Barroso JF. Tunneling nanotubes: a new route for the exchange of components between animal cells. FEBS Lett. 2007;581(11):2194–201.

37. Amano M, Nakayama M, Kaibuchi K. Rho-kinase/ROCK: A key regulator of the cytoskeleton and cell polarity. Cytoskeleton (Hoboken). 2010;67(9):545–54.

38. Engler AJ RL, Wong JY, Picart C, Discher D. Surface probe measurements of the elasticity of sectioned tissue, thin gels and polyelectrolyte multilayer films: correlations between substrate stiffness and cell adhesion. Surface Sci 2004;570(1):11.

39. Del Alamo JC, Meili R, Alonso-Latorre B, Rodriguez-Rodriguez J, Aliseda A, Firtel RA, et al. Spatio-temporal analysis of eukaryotic cell motility by improved force cytometry. Proc Natl Acad Sci U S A. 2007;104(33):13343–8.

40. del Alamo JC, Meili R, Alvarez-Gonzalez B, Alonso-Latorre B, Bastounis E, Firtel R, et al. Three-dimensional quantification of cellular traction forces and mechanosensing of thin substrata by fourier traction force microscopy. PLoS One. 2013;8(9):e69850.

41. Giannotta M, Trani M, Dejana E. VE-cadherin and endothelial adherens junctions: active guardians of vascular integrity. Dev Cell. 2013;26(5):441–54.

42. Serrano R, Aung A, Yeh YT, Varghese S, Lasheras JC, Del Alamo JC. Three-Dimensional Monolayer Stress Microscopy. Biophys J. 2019;117(1):111–28.

43. Beckers CM, Knezevic N, Valent ET, Tauseef M, Krishnan R, Rajendran K, et al. ROCK2 primes the endothelium for vascular hyperpermeability responses by raising baseline junctional tension. Vascul Pharmacol. 2015;70:45–54.

44. Valent ET, van Nieuw Amerongen GP, van Hinsbergh VW, Hordijk PL. Traction force dynamics predict gap formation in activated endothelium. Exp Cell Res. 2016;347(1):161–70.

45. Mebius MM, van Genderen PJ, Urbanus RT, Tielens AG, de Groot PG, van Hellemond JJ. Interference with the host haemostatic system by schistosomes. PLoS Pathog. 2013;9(12):e1003781.

46. Shariati F, Perez-Arellano JL, Carranza C, Lopez-Aban J, Vicente B, Arefi M, et al. Evaluation of the role of angiogenic factors in the pathogenesis of schistosomiasis. Exp Parasitol. 2011;128(1):44–9.

47. Nemethova M, Auinger S, Small JV. Building the actin cytoskeleton: filopodia contribute to the construction of contractile bundles in the lamella. J Cell Biol. 2008;180(6):1233–44.

48. Eugenin EA, Gaskill PJ, Berman JW. Tunneling nanotubes (TNT) are induced by HIV-infection of macrophages: a potential mechanism for intercellular HIV trafficking. Cell Immunol. 2009;254(2):142–8.

49. Abounit S, Zurzolo C. Wiring through tunneling nanotubes--from electrical signals to organelle transfer. J Cell Sci. 2012;125(Pt 5):1089–98.

50. Austefjord MW, Gerdes HH, Wang X. Tunneling nanotubes: Diversity in morphology and structure. Commun Integr Biol. 2014;7(1):e27934.

51. De Pascalis C, Etienne-Manneville S. Single and collective cell migration: the mechanics of adhesions. Mol Biol Cell. 2017;28(14):1833–46.

52. Sherer NM. Long-distance relationships: do membrane nanotubes regulate cell-cell communication and disease progression? Mol Biol Cell. 2013;24(8):1095–8.

53. Gerdes HH, Rustom A, Wang X. Tunneling nanotubes, an emerging intercellular communication route in development. Mech Dev. 2013;130(6–8):381–7.

54. Race GJ, Martin JH, Moore DV, Larsh JE, Jr. Scanning and transmission electronmicroscopy of Schistosoma mansoni eggs, cercariae, and adults. Am J Trop Med Hyg. 1971;20(6):914–24.

55. Sakamoto K, Ishii Y. Fine structure of schistosome eggs as seen through the scanning electron microscope. Am J Trop Med Hyg. 1976;25(6):841–4.

56. Kevin K. Takaki GR, Matthew Berriman, Antonio J. Pagán, and Lalita Ramakrishnan. Schistosoma mansoni Eggs Modulatethe Timing of Granuloma Formationto Promote Transmission. Cell Host & Microbe. 2020;29:10.

57. Baptista AP, Andrade ZA. Angiogenesis and schistosomal granuloma formation. Mem Inst Oswaldo Cruz. 2005;100(2):183–5.

58. Schramm G, Gronow A, Knobloch J, Wippersteg V, Grevelding CG, Galle J, et al. IPSE/alpha-1: a major immunogenic component secreted from Schistosoma mansoni eggs. Mol Biochem Parasitol. 2006;147(1):9–19.

59. Mbanefo EC, Agbo CT, Zhao Y, Lamanna OK, Thai KH, Karinshak SE, et al. IPSE, an abundant egg-secreted protein of the carcinogenic helminth Schistosoma haematobium, promotes proliferation of bladder cancer cells and angiogenesis. Infect Agent Cancer. 2020;15:63.

60. Jourdane J, Theron, A. . Larval development: eggs to cercariae. In: Rollinson D, Simpson AJG, editors. The Biology of Schistosomes: From Genes to Latrines. New York, NY: Academic Press. 1987.

61. Samoil V, Dagenais M, Ganapathy V, Aldridge J, Glebov A, Jardim A, et al. Vesicle-based secretion in schistosomes: Analysis of protein and microRNA (miRNA) content of exosome-like vesicles derived from Schistosoma mansoni. Sci Rep. 2018;8(1):3286.

62. Parsons JT, Horwitz AR, Schwartz MA. Cell adhesion: integrating cytoskeletal dynamics and cellular tension. Nat Rev Mol Cell Biol. 2010;11(9):633–43.

63. Klein EA, Yin L, Kothapalli D, Castagnino P, Byfield FJ, Xu T, et al. Cell-cycle control by physiological matrix elasticity and in vivo tissue stiffening. Curr Biol. 2009;19(18):1511–8.

64. Pinto-Almeida A, Mendes T, de Oliveira RN, Correa Sde A, Allegretti SM, Belo S, et al. Morphological Characteristics of Schistosoma mansoni PZQ-Resistant and - Susceptible Strains Are Different in Presence of Praziquantel. Front Microbiol. 2016;7:594.

65. Cody JJ, Ittiprasert W, Miller AN, Henein L, Mentink-Kane MM, Hsieh MH. The NIH-NIAID Schistosomiasis Resource Center at the Biomedical Research Institute: Molecular Redux. PLoS Negl Trop Dis. 2016;10(10):e0005022.

66. Dalton JP, Day SR, Drew AC, Brindley PJ. A method for the isolation of schistosome eggs and miracidia free of contaminating host tissues. Parasitology. 1997;115 (Pt 1):29–32.

67. Schindelin J, Arganda-Carreras I, Frise E, Kaynig V, Longair M, Pietzsch T, et al. Fiji: an open-source platform for biological-image analysis. Nat Methods. 2012;9(7):676–82.

68. Alvarez-Gonzalez B, Zhang S, Gomez-Gonzalez M, Meili R, Firtel RA, Lasheras JC, et al. Two-Layer Elastographic 3-D Traction Force Microscopy. Sci Rep. 2017;7:39315.

