## Supplemental Information for "Biomechanical interactions of *Schistosoma mansoni* eggs with vascular endothelial cells facilitate egg extravasation"

| | Mature egg | Immature egg |
|---|---|---|
| Long axis (mm) | 132.7 ± 3.4 | 112.6 ± 2.4 *** |
| Short axis (mm) | 59.7 ± 1.8 | 37.2 ± 1.2 *** |
| Height (mm) | 58.4 ± 1.0 | 37.2 ± 1.2 *** |
| Volume (x$10^4$ $mm^3$) | 16.3 ± 3.0 | 7.6 ± 1.7 * |
| Mean stiffness (MPa) | 9.1 | 4.2 |

**Supplemental Table1. Mature and immature *S. mansoni* egg properties.** To measure the long and short axis, and height, 11 each of mature and immature eggs were used. For each parameter, the comparison between mature and immature eggs was significant: ***, $p<0.001$ by Student's *t*-test. To measure volume, seven eggs were used, and the comparison between mature and immature eggs was significant: * $p<0.05$ by Student's *t*-test. To measure egg stiffness, two eggs in each case were probed by Atomic Force Microscopy.

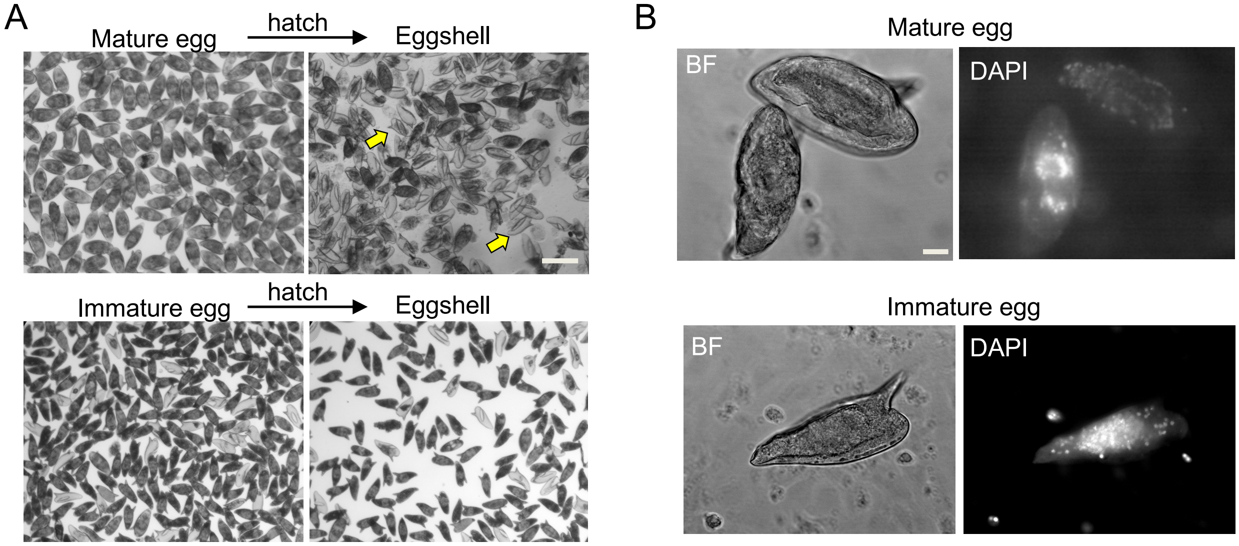


**Supplemental Figure 1. Morphology, hatching ability and staining of mature and immature *S. mansoni* egg embryos. (A)** Mature and immature eggs were cultured for 24 h and then hatching attempted in distilled water under a bright light for 40 min. Unlike mature eggs, of which ~80% had hatched (empty eggshells indicated by arrowheads), immature eggs cannot hatch. Scale bar, 200 μm. All eggs in the experiments were not hatched. **(B)** Images of mature and immature eggs: DAPI was used to stain egg embryo nuclei. Scale bar, 20 μm.

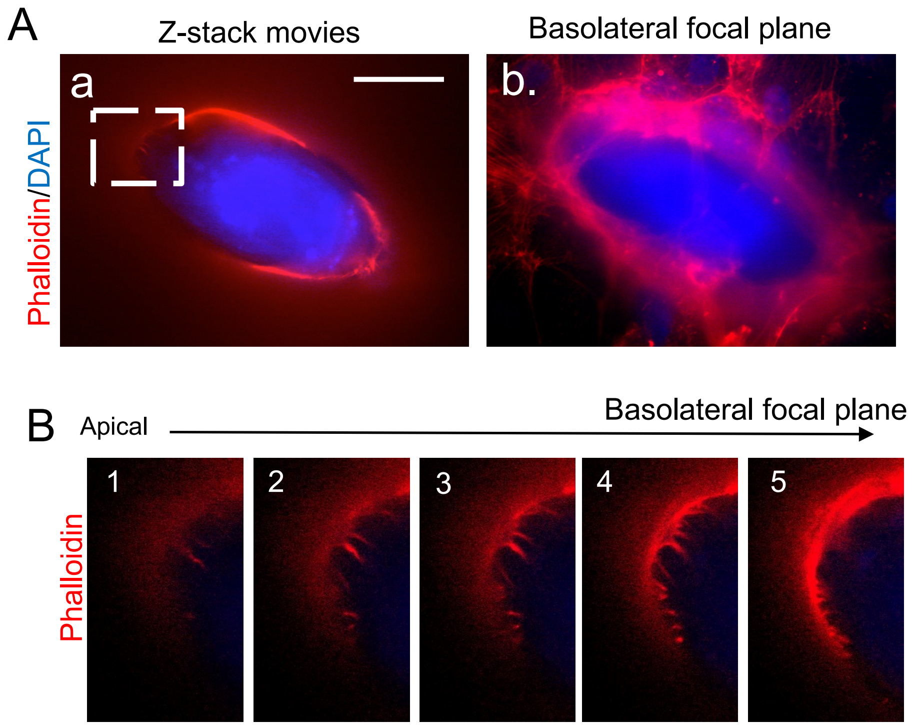


**Supplemental Figure 2. VEC filopodia contain F-actin filaments. (A)** The *S. mansoni* egg was encapsulated by VECs at 4 h. (a) Z-stack images (See SI movie 1) of the distribution of F-actin from the apical to the basolateral focal plane. (b) The basolateral image of the stack images. Scale bar = 50 µm. **(B)** Area of interest, indicated by the white dashed line box in (**A**), enlarged with its component Z-stack sections displaying the stained VEC filopodia.

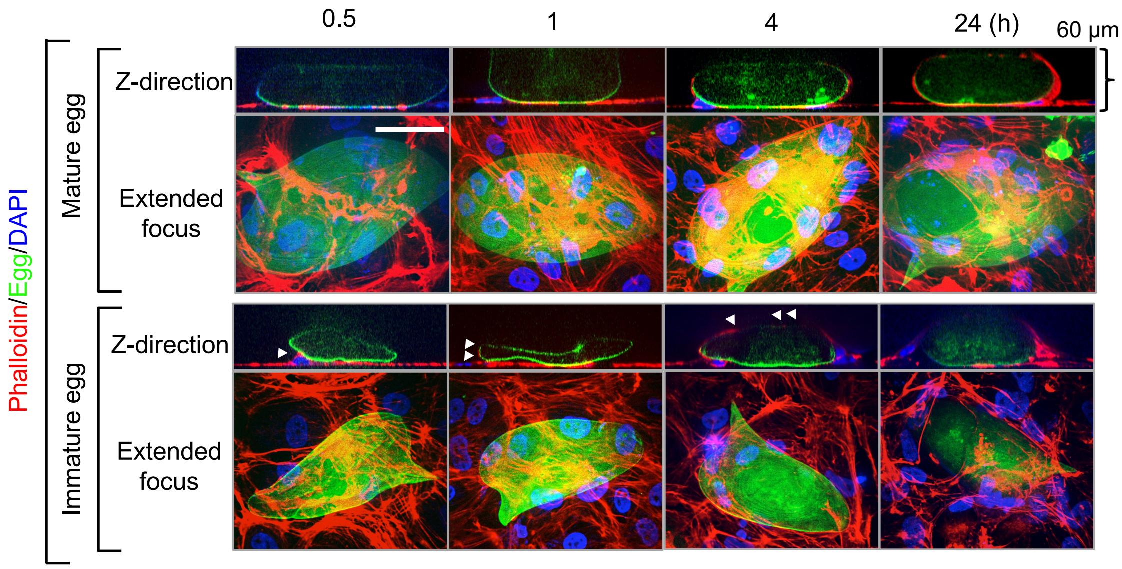


**Supplemental Figure 3. The kinetics of VEC encapsulation of *S. mansoni* eggs.** Representative 3D images of the interaction between mature and immature eggs during the VEC encapsulation process. VECs were immunostained by Phalloidin (red), eggs were auto-fluorescently green, and cell and egg nuclei were immunostained by DAPI. Extended focus images combine the in-focus parts of each image together to produce a single image with an increased depth of field. Scale bar = 50 µm. Z-direction images show the progress of VEC coverage of eggs. The total height is 60 µm. White arrowheads indicate VECs migrating on the side and over the eggs.

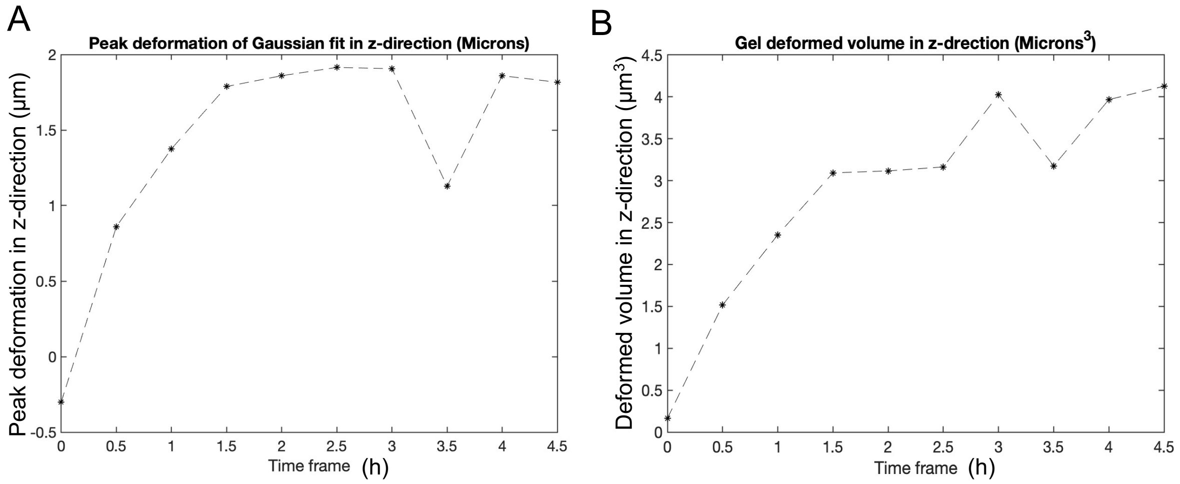


**Supplemental Figure 4. Quantification of peak deformation and gel deformed volume during encapsulation of a live mature *S. mansoni* egg.** To obtain the quantification result for **Fig. 5B**, a cropped area based on peak z-direction displacements was selected at each time point as shown in the red dashed box (**Fig. 5B**) to fit to a 2D Gaussian curve. The peak deformation and deformed gel volume obtained in the z-direction are relative to time point 0.

.

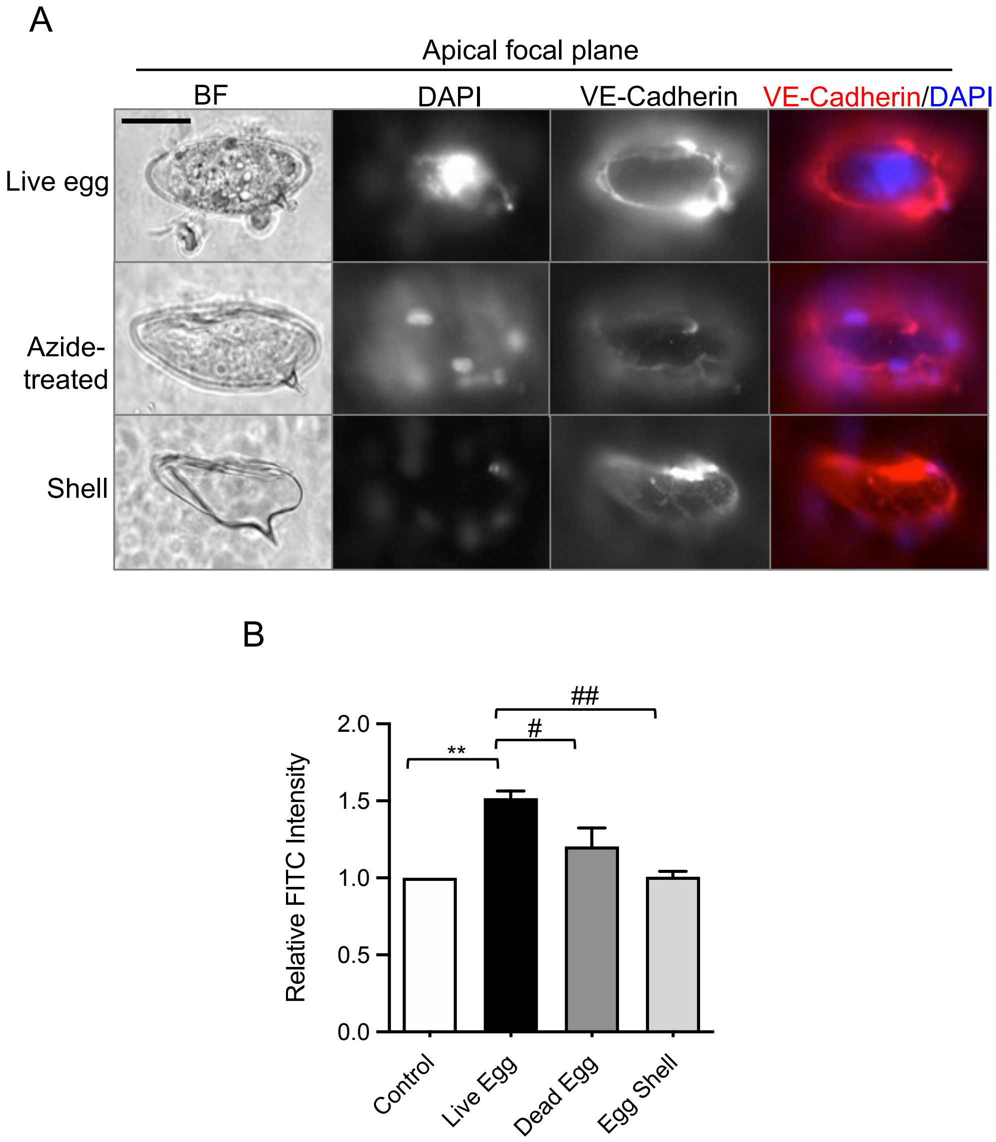


**Supplemental Figure 5. VEC permeability as a function of *S. mansoni* egg vitality. (A)** Live and dead (treated with sodium azide) mature eggs, and eggshells only were incubated with VEC monolayers for 24 h. Apical bright field images were taken. VE-Cadherin (red) and DAPI (blue) were immunostained in the VECs present on the eggshell surface and show how the cells can encapsulate the three egg preparations after 24 h. Scale bar, 50 μm. **(B)** Permeability response of VEC junctions after interaction with live or dead eggs, or eggshells for 4 h. The control condition is VECs monolayers without eggs. Data represent the mean ± s.e.m. **P<0.01, # P<0.05 and ## P<0.01 using the one-way ANOVA with Tukey's multiple comparison test. Three independent experiments were performed.

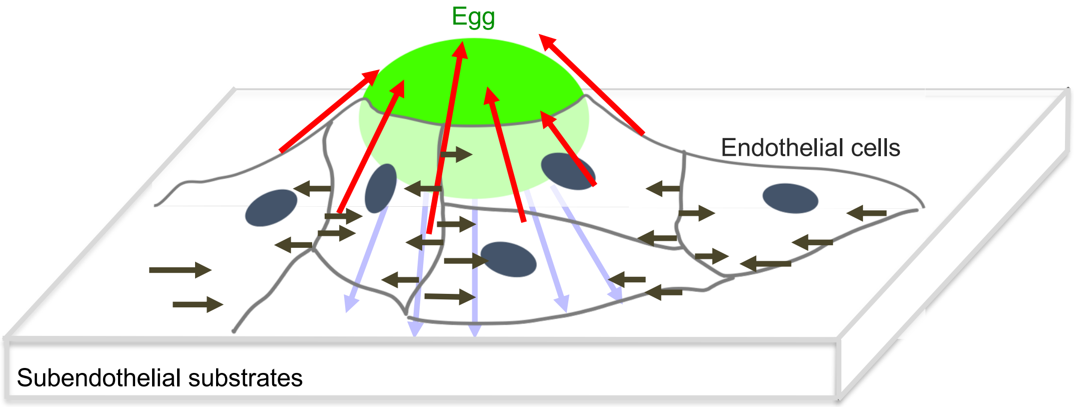


**Supplemental Figure 6. Schematic of the tangential and perpendicular stresses during encapsulation of a *S. mansoni* egg.** Endothelial cells encapsulating a *S. mansoni* egg (green oval) collectively generate pulling upward and inward forces (indicated by red arrows) onto the subendothelial substrate (e.g., basement membrane). These forces are exerted as the leading edge of the endothelium climbs up and over the egg and are required to balance the downward pushing forces that endothelial cells exert on the egg. The downward pushing forces are transmitted to the substrate under the egg's central region (purple arrows). In comparison, the horizontal forces generated by contraction of individual cells contribute a more disorganized tangential stress pattern (black arrows). Due to the downward forces exerted by the endothelial cells that overlie the egg, the integrity of the endothelial monolayer directly underneath the egg is compromised, eventually breaking the cell-cell junctions. *In vivo*, this process would force the egg into the underlying tunica layers of the blood vessel, thus initiating the migration of the egg away from the blood vessel.

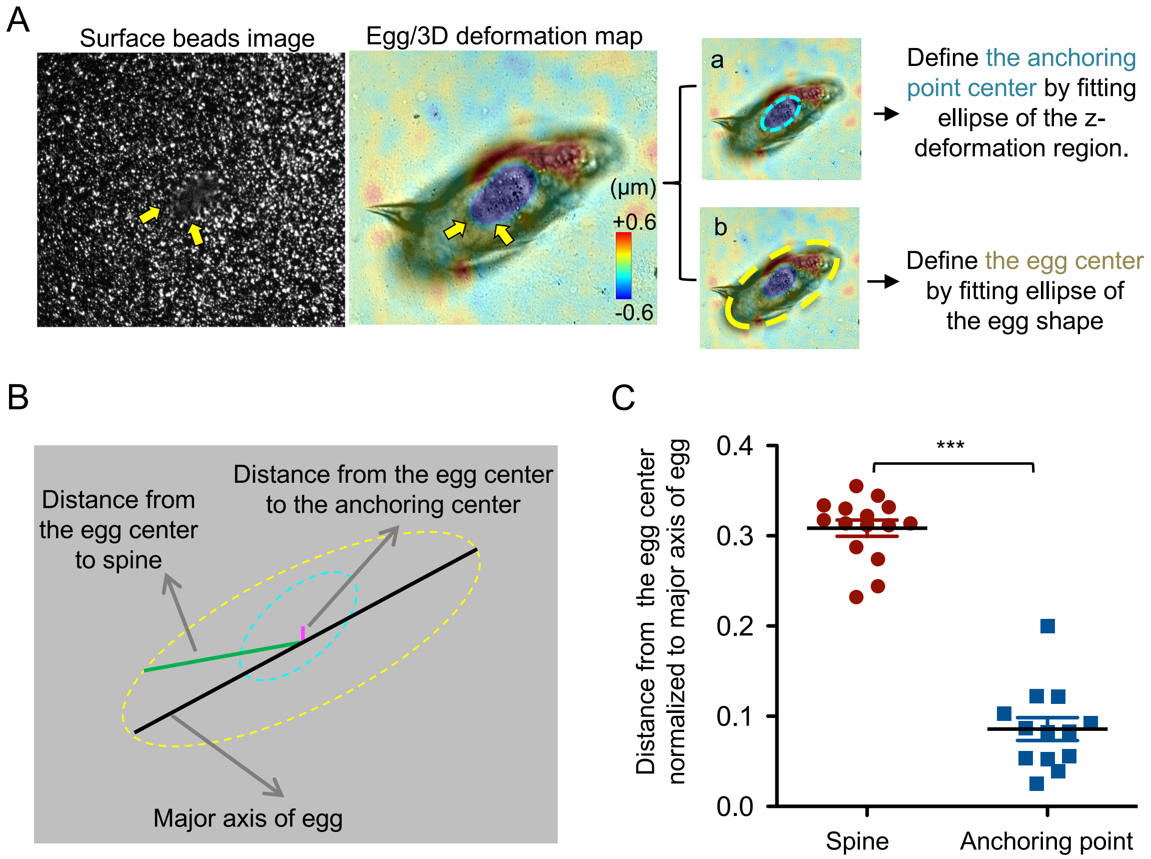


**Supplemental Figure 7. Analysis of the spatial localization of the *S. mansoni* egg's lateral spine and the anchoring point relative to the egg basal centroid. (A)** As introduced in **Fig. 4A** and **4B** , we defined the center of the anchoring point by fitting the ellipse shape for the z-deformation region (a). In addition, we define the egg center by fitting the ellipse shape for the egg shape (b). Scale bar, 50 µm. **(B)** The distance from the egg center to the egg's lateral spine (green line) and from the egg center to the anchoring point (pink) were calculated by using the FIJI imaging processing software. The major axis of the egg was used to normalize for the variation in the size of each egg. **(C)** Quantification of the distance from the egg's center to the lateral spine and to the anchoring point. Data were normalized to the length of the long axis of each egg. For each condition, 12-15 eggs were used: ****, $p<0.0001$ by Student's *t*-test.
